# Opto-MDMi: a dual-lock optogenetic system for robust activation of endogenous p53

**DOI:** 10.64898/2026.04.13.718310

**Authors:** Tatsuki Tsuruoka, Takashi Sumikama, Sachiya Nakashima, Yuhei Goto, Kazuhiro Aoki

## Abstract

Optogenetics has emerged as a powerful technology for manipulating biological functions with high spatiotemporal resolution, yet the precise control of endogenous molecules remains a significant challenge. In this study, we developed Opto-MDMi, a dual-lock optogenetic platform designed to control the activity of endogenous p53, a master regulator of cell cycle and apoptosis. The p53 pathway is strictly governed by its negative regulators, MDM2 and MDMX, which inhibit p53 through direct binding and ubiquitination. Our system integrates two distinct light-responsive modules: Opto-MDMi (LOVTRAP), which regulates the nuclear translocation of p53-activating peptides, and Opto-MDMi (LOV2-PMI), which controls the binding activity of these peptides by photocaging them within the AsLOV2 domain. Through extensive *in vitro* screening and live-cell assays, we discovered that truncating the Jα helix of LOV2 effectively restricts the movement of fused inhibitory peptides, thereby masking their interaction with MDM2/MDMX under dark conditions. By combining these two regulatory layers into a dual-lock system, we achieved robust light-dependent activation of endogenous p53 while significantly suppressing basal activity in the dark. Our findings not only provide a potent tool for p53 research but also establish a general design principle for optogenetically regulating functional peptides with the LOV2 domain, offering a versatile framework for the future development of optogenetic actuators.

## Introduction

Optogenetics has become an essential tool for manipulating biological functions with light. This concept was initially demonstrated in the neuroscience field using light-responsive ion channels and pumps (Yizhar et al., 2011). Subsequently, diverse photosensory proteins have been adapted as optogenetic tools, leading to their widespread applications across broad fields of life sciences (Repina et al., 2017). In particular, the development of optogenetic tools targeting endogenous molecules has advanced significantly over the past decade; numerous optogenetic tools have been reported that enable the control of intermolecular interactions and the manipulation of signaling pathways (Manoilov et al., 2021). These tools are typically composed of light-responsive domains of photosensory proteins derived from plants, fungi, or bacteria, combined with various functional effector proteins. The conformational change triggered by light depends heavily on the specific light-responsive domain employed. Consequently, strategies for developing optogenetic tools vary according to the light-responsive domain.

Among photosensory domains, the light-responsive LOV2 domain of *Avena Sativa* phototropin1 (AsLOV2) is widely used as the backbone for designing a broad range of optogenetic tools (Hoffmann et al., 2018). AsLOV2 is a relatively small, single-component domain of approximately 140 residues, featuring a C-terminal amphipathic helix known as the Jα helix (Harper et al., 2003). Upon blue light illumination, the flavin mononucleotide (FMN) chromophore forms a covalent bond with a cysteine residue within LOV2, triggering the unfolding of the Jα helix (Harper et al., 2003; Halavaty and Moffat, 2007; Zayner et al., 2012). Leveraging this substantial conformational change enables us to design many types of optogenetic tools for light-mediated manipulation of small GTPase, tyrosine kinase/phosphatase, phospholipase, and transcription factor, to name a few (Wu et al., 2009; Dagliyan et al., 2016; Hongdusit et al., 2020; Zhu et al., 2023; Li et al., 2024). Furthermore, sophisticated platforms have been established to convert the conformational change in LOV2 into light-dependent protein-protein interactions (Wang et al., 2016; Guntas et al., 2015; Strickland et al., 2012). For example, an optogenetic system called LOVTRAP, which induces light-dependent dissociation between proteins, was developed by screening of fragments that bind selectively to the structure of the LOV2 domain under dark conditions (Wang et al., 2016).

Another promising strategy for developing LOV2-based optogenetic tools involves photocaging effector peptides or small domains onto the C-terminal Jα helix (Baarlink et al., 2013; He et al., 2015; Paonessa et al., 2016; Niopek et al., 2014, 2016; Yumerefendi et al., 2015, 2016; Lungu et al., 2012; Yi et al., 2014; Murakoshi et al., 2017; Melero-Fernandez de Mera et al., 2017; Strickland et al., 2012). In particular, optogenetic systems that achieve light-dependent nuclear import and export by embedding nuclear localization/export signal sequences within Jα-helix have become a versatile platform for controlling protein subcellular localization (Yumerefendi et al., 2015, 2016; Niopek et al., 2014, 2016). Compared to inserting LOV2 directly into a target protein, this approach offers a relatively easy way to obtain LOV2-based light-responsive modules. Especially for functional peptides, it is possible to utilize not only naturally existing amino acid sequences but also those identified through established screening methods, such as phage or mRNA display (Newton et al., 2020; Jaroszewicz et al., 2022). Additionally, previously reported functional peptides are accessible through public databases (Quiroz et al., 2021). These advantages make the incorporation of peptides into LOV2 a highly promising strategy for rapid generation of optogenetic tools. However, despite multiple reports to date, general design principles for caging peptides in the LOV2 domain remain poorly defined.

In this study, we developed Opto-MDMi, a tool capable of light-dependent activation of endogenous p53, by employing an inhibitory peptide targeting MDM2/MDMX, which are negative regulators of p53. Opto-MDMi has two components: Opto-MDMi (LOVTRAP) and Opto-MDMi (LOV2-PMI). Opto-MDMi (LOVTRAP) enables the targeted delivery of p53-activating modules into the nucleus by using the LOVTRAP light-responsive dissociation system (Wang et al., 2016). Meanwhile, Opto-MDMi (LOV2-PMI) facilitates the light-dependent activation of effector peptides by caging the p53 activation peptides within LOV2. Finally, by combining these systems, we successfully induced light-dependent activation of endogenous p53 while maintaining its low basal level under dark conditions. The workflow we employed—combining *in vitro* screening and live-cell assays in the development of Opto-MDMi (LOV2-PMI)—is broadly applicable to the development and evaluation of other LOV2-photocaged peptide modules. Furthermore, the insights obtained in this study regarding photocaging peptides to LOV2 may serve as general guidelines for the future design of similar optogenetic tools.

## Results

### Light-dependent translocation of the inhibitory peptide for p53-MDM2/MDMX interactions

Since the p53 transcription factor functions primarily in the nucleus, we first investigated whether spatial manipulation of peptides that activate the p53 pathway could control p53 activity. The transcriptional activity of p53 is strictly regulated by its negative regulators, MDM2 and MDMX (Fig. 1A) (Oliner et al., 1993; Wade et al., 2013; Klein et al., 2021). MDM2 promotes p53 ubiquitination and subsequent degradation, thereby suppressing its expression level (Kubbutat et al., 1997; Haupt et al., 1997; Honda et al., 1997). Additionally, both MDM2 and MDMX inhibit p53 activity by directly binding to p53 and masking its transcriptional activation domain (Oliner et al., 1993; Shvarts et al., 1996). This interaction is part of a negative feedback loop, as MDM2 and MDMX are themselves direct target genes of p53 (Zauberman et al., 1995; Phillips et al., 2010). It has been reported that peptides such as PMI (p53-MDM2/MDMX inhibitor) or PMI-M3 enable activation of the p53 pathway by effectively inhibiting these interactions between p53 and MDM2/MDMX (Fig. 1B) (Pazgier et al., 2009; Li et al., 2021). Given that these peptide sequences can be genetically encoded and possess distinct *in vitro* affinities for MDM2 and MDMX, we adopted them for development of our optogenetic tools.

**Figure 1.**
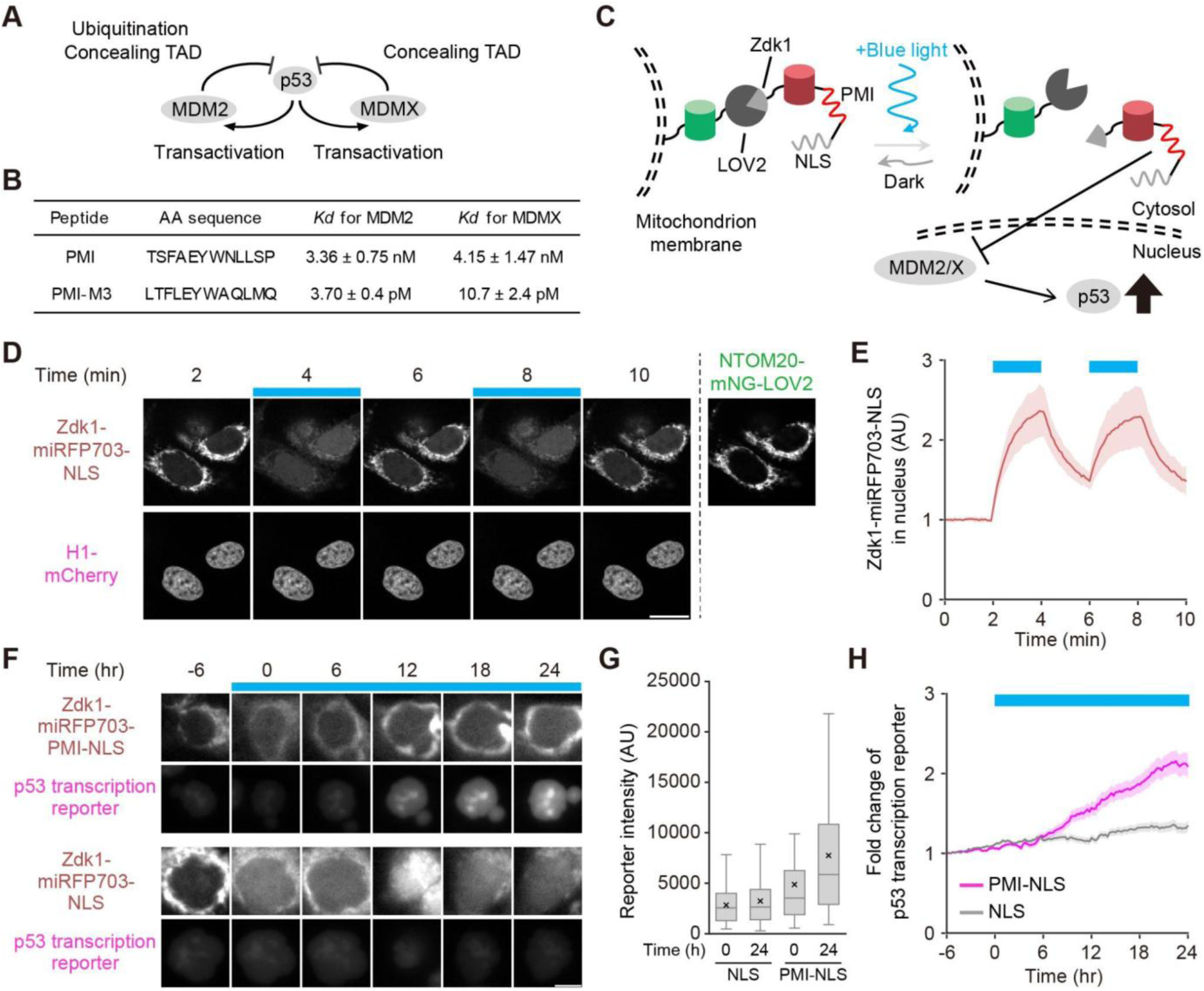
Design and evaluation of the Opto-MDMi (LOVTRAP) system. A. Schematic of the p53-MDM2/MDMX negative feedback loop. B. Amino acid sequence and *in vitro K*_d_ value of PMI peptides for MDM2 and MDMX. C. Schematic illustration of the Opto-MDMi (LOVTRAP) system. D. Light-dependent nuclear accumulation of the Opto-MDMi (LOVTRAP) actuator. Blue boxes indicate periods of blue light illumination. H1-mCherry serves as a nuclear marker. Scale bar, 20 µm. E. Quantification of the actuator levels in the nucleus. Data are presented as mean ± s.d. (n = 10 cells). F. Representative images showing changes in actuator localization and p53 transcription reporter signal. Scale bar, 10 µm. G. Distribution of p53 reporter fluorescence intensity at 0 and 24 hours from the start of blue light illumination. Horizontal lines and crosses indicate the medians and means, respectively. Boxes represent the 25th and 75th percentiles; whiskers indicate the range between the maximum and minimum values excluding outliers. H. Temporal changes of the p53 transcription reporter in each cell line. Data represent mean ± s.e.m. (PMI-NLS, n = 122 cells; NLS, n = 125 cells).

Although several single-component light-dependent translocation systems have been reported to control the intracellular localization of a protein of interest (POI) (Niopek et al., 2014, 2016; Yumerefendi et al., 2015, 2016), a previous attempt similar to ours failed to induce efficient nuclear translocation of PMI peptide modules due to the limitations of the light-dependent nuclear import system (Wehler and Di Ventura, 2019). To overcome this issue, we employed the LOVTRAP light-dependent dissociation system to achieve more robust nuclear translocation of the PMI peptide module (Wang et al., 2016). LOVTRAP consists of the photosensory domain LOV2 and its binding partner, Zdk1. Blue light illumination triggers a conformational change in LOV2, causing the dissociation of Zdk1 from LOV2. Hence, by anchoring either LOV2 or Zdk1 to the mitochondrial outer membrane via NTOM20 and fusing the other to the POI, the POI can be sequestered at the mitochondria in the dark and released into the cytoplasm upon blue light illumination.

Based on this principle, we designed an optogenetic system named Opto-MDMi (LOVTRAP) (Fig. 1C and Fig. S1A). In this system, Zdk1-miRFP703-PMI-NLS (the “actuator”) binds to mitochondria-localized NTOM20-mNeonGreen-LOV2 (the “localizer”) under dark conditions (Fig. 1C, left). Upon blue light illumination, the actuator dissociates from the localizer and translocates to the nucleus via the nuclear localization sequence, NLS (Fig. 1C, right), where it is expected to induce p53 activation by inhibiting p53-MDM2/MDMX interaction. We confirmed that blue light illumination efficiently induced nuclear translocation of the actuator in HeLa cells (Fig. 1D, E and Movie 1). Notably, using a stronger NLS (3xNLS) resulted in substantial nuclear localization even in the dark (Fig. S1B, C), indicating that the efficiency of light-induced nuclear translocation depends on the balance between the NLS strength and the LOVTRAP system.

Next, we established HCT116 cell lines (p53 wild-type) stably expressing a p53 transcription reporter, the Opto-MDMi (LOVTRAP) localizer, and its actuators to confirm whether our system could activate endogenous p53 via light stimulation. The p53 transcription reporter was previously developed in our laboratory, featuring the promoter region with high specificity for p53 (Tsuruoka et al., 2023, 2025). We verified that the p53 activation—induced by 24 hours treatment with etoposide (Nitiss, 2009; Yang et al., 2018) or nutlin-3a (Vassilev et al., 2004)—increases the reporter signal in a dose-dependent manner (Fig. S1D and E). To minimize leakage of the actuator into the nucleus under dark conditions, we enriched the cell population with high localizer expression via cell sorting and used them for subsequent experiments. This cell population exhibited transcriptional activation of p53 upon blue light illumination (Fig. 1F, upper panels, Fig. 1G, and Movie 2, left). In contrast, negative control cells lacking the PMI peptide showed almost no p53 activation (Fig. 1F, lower panels, Fig. 1G, and Movie 2, right). We should note that cells expressing the PMI-NLS actuator displayed relatively higher basal reporter activity than negative control, suggesting some leakage of PMI activity under dark conditions (Fig. 1G). Quantification revealed that Opto-MDMi (LOVTRAP)-expressing cells showed, on average, a 2.1-fold increase in p53 transcription reporter activity after 24 hours of blue light illumination (Fig. 1H). However, prior to cell sorting, we observed that the expression level of the PMI-containing actuator was significantly lower than that of control, with its distribution largely overlapping that of the parental (non-expressing) cells (Fig. S1F). Furthermore, when attempting to establish stable cell lines using the PMI-M3 peptide, which possesses even stronger inhibitory activity than PMI, we were unable to obtain a cell population with fluorescence levels detectable by fluorescence microscopy. Based on these results and the elevated basal reporter activity, we concluded that the persistent activity of the PMI/PMI-M3 peptides even under dark conditions—either by constitutively activating p53 or by inducing actuator degradation through MDM2 interaction—hampers the performance of this optogenetic system. For this reason, we next focused on improving the regulatory control of the actuator fragment.

### *in vitro* screening of the light-responsive LOV2-PMI inhibitory modules

We explored a strategy of caging PMI peptides within the LOV2 domain to enable light-dependent control over the interaction between MDM2/MDMX and the PMI peptide (Fig. 2A). To obtain actuator fragments with robust light responsiveness, we established an *in vitro* binding assay based on previous reports (Fig. 2B) (Melero-Fernandez de Mera et al., 2017). The p53-binding domains of MDM2 and MDMX were expressed and purified as GST fusion proteins in *E. coli* and immobilized onto beads as the bait (Fig. 2C). The actuator fragments (prey) were also expressed in *E. coli* and prepared as total lysate. For these fragments, we utilized LOV2 mutants that mimic either the dark-state (*dsm*; C450A) or light-state (*lsm*; I539E) conformation (Yi et al., 2014; Melero-Fernandez de Mera et al., 2017). Bait and prey fractions were mixed, and candidates of light-responsive LOV-PMI fragments were screened on the basis of their binding activity to the p53 binding domain of MDM2/MDMX.

**Figure 2.**
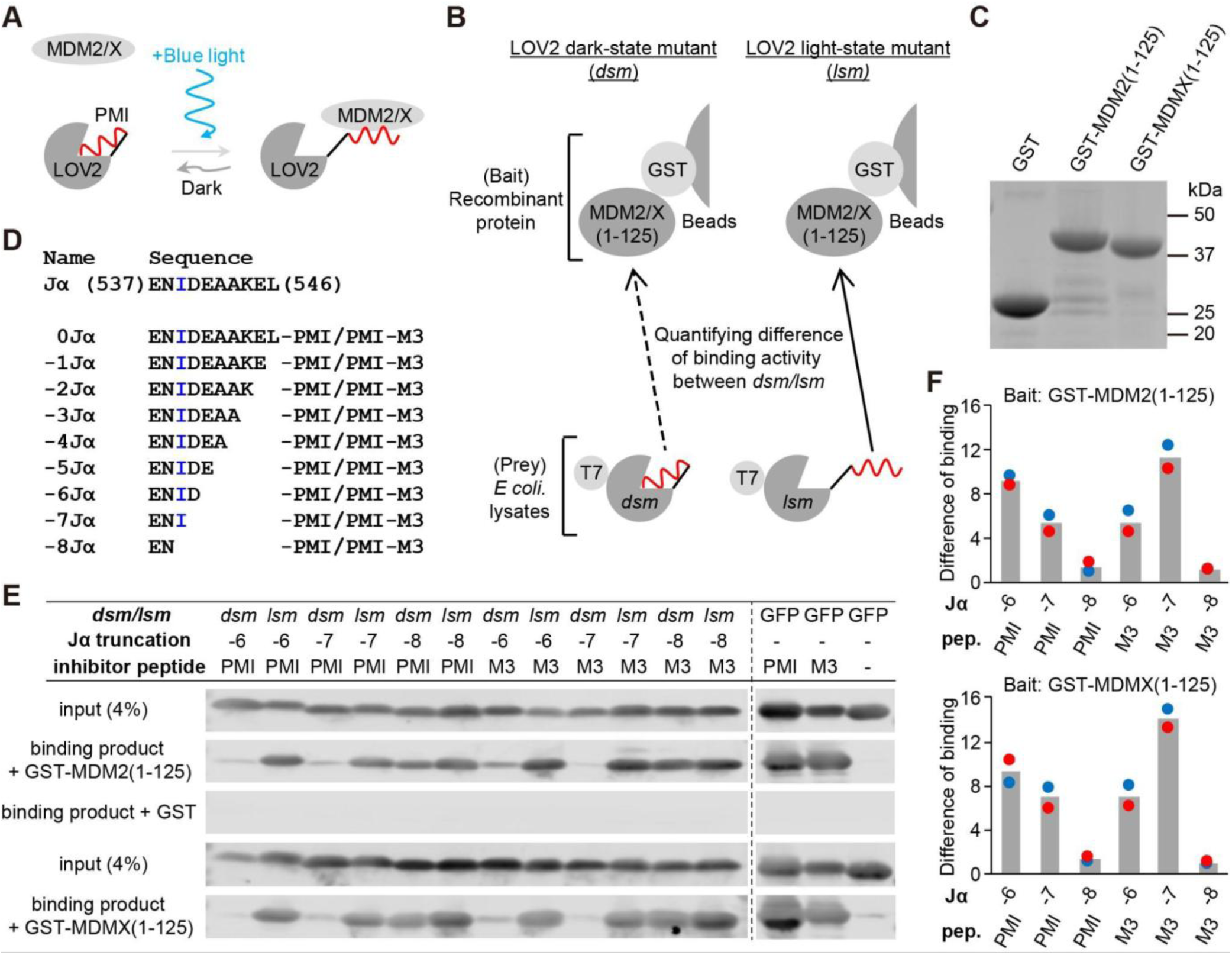
*in vitro* screening of light-responsive inhibitory modules. A. Schematic of the light-responsive LOV2-PMI chimera module. B. Workflow of the *in vitro* binding assay. T7 denotes the T7 tag used for detection. C. Representative CBB-stained SDS-PAGE gel of purified recombinant bait protein. D. Library of the LOV2 Jα-helix truncation mutants fused with PMI or PMI-M3 peptides. The critical residue Ile539 is highlighted in blue. E. Representative western blot detecting LOV2-PMI/PMI-M3 fragments bound to GST-MDM2(1–125) or GST-MDMX(1–125). The LOV2(–8Jα)-PMI/PMI-M3 fragment, truncated up to I539, serves as a control where light responsiveness is expected to be lost. GST is a negative control bait fragment. GFP-PMI/PMI-M3 and GFP are positive and negative control prey fragments, respectively. F. Differences in binding activity between *dsm* or *lsm* of each LOV2-PMI truncation mutant to GST-MDM2(1–125) (upper plot) or GST-MDMX(1–125) (lower plot). Blue and red points indicate experimental replicates.

While previous studies have developed light-responsive LOV2-peptide chimeras by using sequence homology between the Jα helix and effector peptides (Lungu et al., 2012; Niopek et al., 2014, 2016; Yumerefendi et al., 2015, 2016), no clear sequence homology was found between the Jα helix and the PMI peptide. Therefore, we first tested N- or C-terminal truncations of the peptide itself, as previously demonstrated (Melero-Fernandez de Mera et al., 2017). An expression library was generated by fusing either the truncated PMI or PMI-M3 peptide to the C-terminus of the intact LOV2 domain (Fig. S2A). Using this expression library, we found that the binding activity to the p53 binding domain of MDM2 was reduced depending on the truncated number of amino acid residues in the PMI/PMI-M3 peptide regardless of *dsm/lsm* (Fig. S2B, C). However, no substantial difference between *dsm* and *lsm* was observed in the binding activity in almost all variants (Fig. S2D). The only exception was the PMI-M3 mutant with C-terminal three residue truncation, which exhibited about a 5-fold difference in binding activity between *dsm* and *lsm*. Nevertheless, we reasoned that this variant would likely be ineffective in living cells, due to its decreased absolute binding activity (Fig. S2B, C).

It has been reported that truncating the amino acids on the Jα helix side potentially could make the LOV2-peptide chimera module responsive to light (Strickland et al., 2012). Hence, we next investigated the effect of truncating the Jα helix itself. Since the Ile539 mutation constitutively undocks the Jα helix, we constructed a library of fusion fragments consisting of PMI/PMI-M3 and LOV2 truncated up to Ile539 (Fig. 2D). Screening revealed several LOV2-PMI/PMI-M3 fragments with marked differences in binding activity to the p53 binding domain of MDM2/MDMX between *dsm* and *lsm* (Fig. 2E, F; all screening data are in Fig. S2E, F). When comparing the p53 binding domains of MDM2 and MDMX used as baits, a similar trend in binding activity was observed across the examined LOV2-PMI variants (see upper and lower panels of Fig. 2E). In particular, LOV2(−6Jα)-PMI and LOV2(−7Jα)-M3 exhibited the most significant light-dependent changes and were therefore selected for further characterization.

### Interaction between LOV2-PMI modules and p53 binding domain of MDM2 in cultured cells

We developed a translocation assay system to confirm whether the LOV2-PMI module obtained through *in vitro* screening interacts with MDM2 in a light-dependent manner in cultured cells (Fig. 3A). In this system, the p53 binding domain of MDM2 was anchored to the plasma membrane via fusion with the C-terminal domain of HRas (HRasCT), while the LOV2-PMI modules were expressed in the cytoplasm. We then monitored the translocation of LOV2-PMI modules to the plasma membrane upon blue light illumination. Indeed, the LOV2(–6Jα)-PMI and LOV2(–7Jα)-M3 fragments demonstrated rapid and reversible translocation to the plasma membrane in a blue light-dependent manner (Fig. 3B, second and third rows, Fig. 3C, and Movie 3, center left and center right). On the other hand, LOV2 alone showed no change in localization with or without blue light exposure (Fig. 3B, first row Fig. 3C, and Movie 3, far left). Furthermore, the LOV2(–8Jα)-PMI-M3 fragment, which was expected to lack light responsiveness due to Jα helix truncation up to I539, was constitutively localized to the plasma membrane even under dark conditions (Fig. 3B, fourth row and Movie 3, far right). These results indicate that the LOV2-PMI module screened by the *in vitro* binding assay interacts with MDM2 in a light-dependent manner in living cells. Given the high sequence homology between MDM2 and MDMX and their similar binding modes, it is inferred that MDMX also interacts in a light-dependent manner within cells.

**Figure 3.**
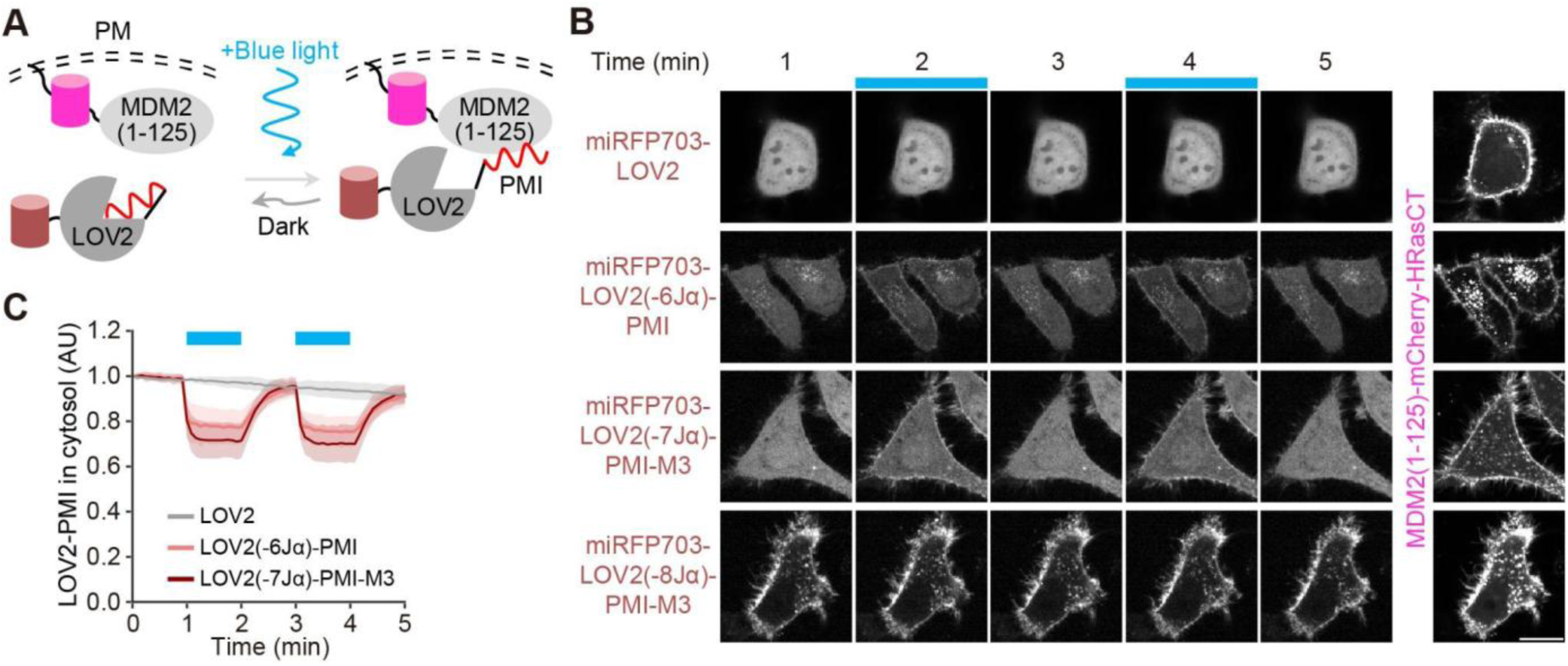
Light-dependent interaction between LOV2-PMI fragments and MDM2 in cultured cells. A. Schematic illustration of the live cell-based translocation assay. B. Light-dependent changes in localization of the Opto-MDMi (LOV2-PMI) actuator fragment identified through *in vitro* screening. Blue boxes indicate the time points at which light illumination was applied. Scale bar, 20 µm. C. Quantification of the Opto-MDMi (LOV2-PMI) actuator cytosolic level. Data are presented as mean ± s.d. (LOV2, n = 15 cells; LOV2(–6Jα)-PMI, n = 30 cells; LOV2(–7Jα)-PMI-M3, n = 20 cells).

When performing long-term live imaging with optogenetic systems, phototoxicity sometimes can be a serious concern. The wild-type LOV2 domain possesses relatively fast off-kinetics (approximately several tens of seconds) among common photoreceptor domains, which necessitates continuous light illumination to maintain its conformational change induced by light (Salomon et al., 2000). Therefore, reducing light intensity or decreasing illumination frequency could minimize phototoxicity. Since a LOV2(V416L) mutant, which has slower off-kinetics than wild-type LOV2 (Kawano et al., 2013), is already known, we carried out an *in vitro* binding assay using this mutant. We found that several LOV2(V416L)-PMI modules also showed differences in binding activity to the p53 binding domain of MDM2 between *dsm* and *lsm* that was comparable to or greater than that of wild-type LOV2 (Fig. S3A, B). Surprisingly, in the subsequent translocation assay in living cells, the majority of the LOV2(V416L)-PMI module remained localized to the plasma membrane even under dark conditions, and only a small fraction of the LOV2(V416L)-PMI module exhibited blue-light-dependent change in localization (Fig. S3C, D). Because the dark state of the LOV2(V416L)-PMI fragment probably retains partial binding activity to MDM2, it is not suitable for the use in optogenetic regulation of p53. The reason for the discrepancy between *in vitro* binding assay and cultured cell-based translocation assay for the LOV2(V416L) mutant remains unclear at present. However, these results highlight the importance of verifying interactions not only *in vitro* but also within live cells.

### Mechanism of light responsivity predicted by molecular dynamics simulation

To understand the mechanism of light-dependent interaction between the LOV2-PMI modules and MDM2/MDMX, we compared the predicted structures of each fusion fragment using AlphaFold 3 (Fig. 4A and Movie 4) (Abramson et al., 2024). The prediction results suggested that PMI/PMI-M3 peptides protruded from the C-terminus of the full-length LOV2 (Fig. 4A, first and third panels).

**Figure 4.**
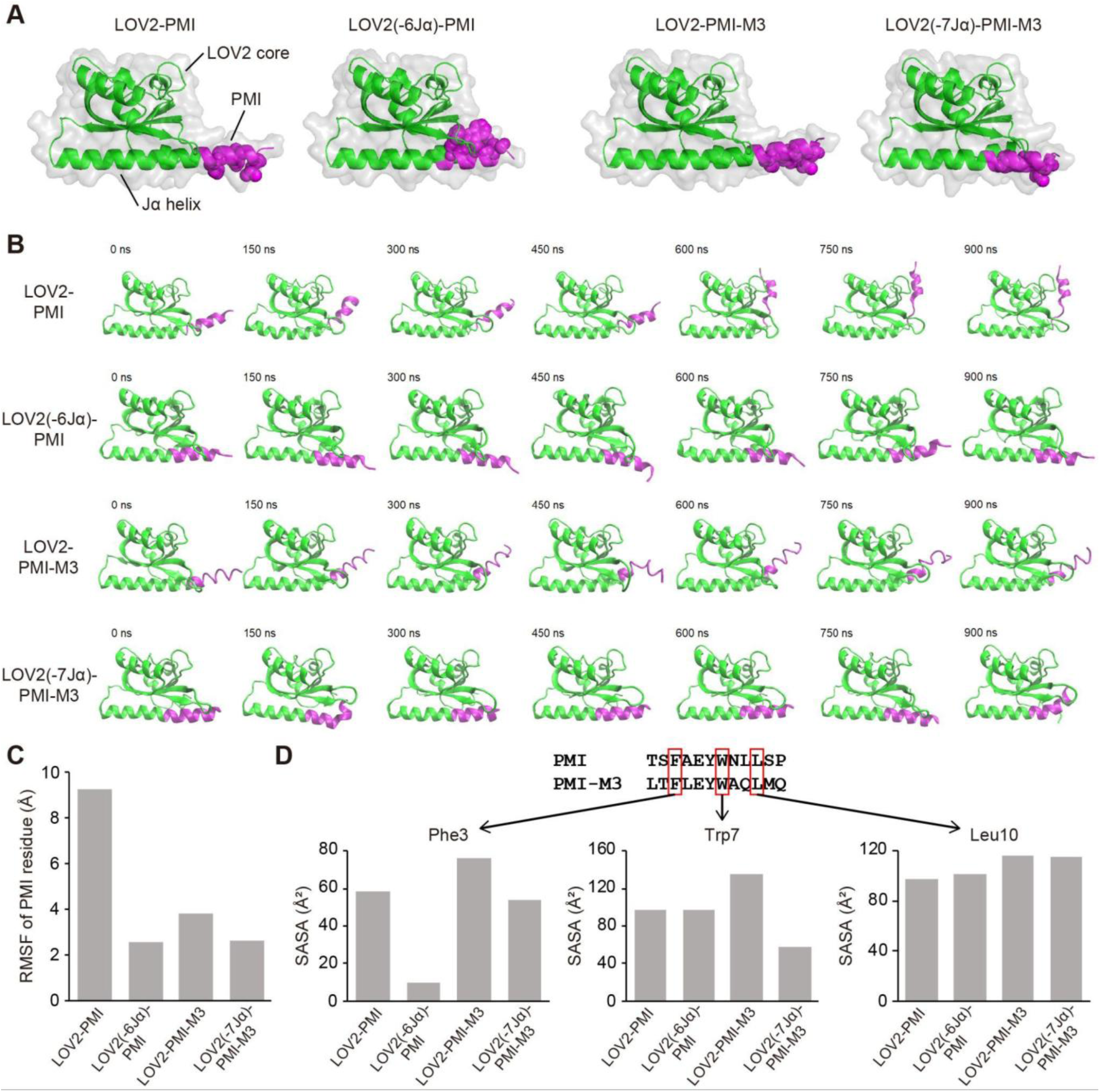
Proposed mechanism of light-dependent interaction revealed by molecular dynamics simulations. A. Protein structures of screened LOV2-PMI fragments predicted by AlphaFold 3. The amino acid residues drawn in light green and magenta indicate the LOV2 core domain and the PMI peptide, respectively. The Phe3, Trp7, and Leu10 residues in PMI/PMI-M3 peptides, known to be particularly important for interactions with MDM2/MDMX, were displayed in the ball-and-stick model to highlight their side chains. B. Snapshots of the Opto-MDMi (LOV2-PMI) protein structure obtained by molecular dynamics (MD) simulation. Protein structures were displayed every 150 nanoseconds based on the results of the MD simulation. C. Calculated root mean square fluctuation (RMSF) value of the Cα atom in the PMI/PMI-M3 peptides. D. Calculated solvent accessible surface area (SASA) value for the Phe3, Trp7, and Leu10 residues in PMI/PMI-M3 peptides.

Furthermore, focusing on residues crucial for interaction with MDM2/MDMX (Phe3, Trp7, Leu10; common to PMI/PMI-M3), all these residues were predicted to be exposed on the protein surface. In contrast, in the fusion fragments with the Jα helix-truncated LOV2 obtained from screening, PMI/PMI-M3 was positioned relatively internally within the protein, and the residues critical for interaction with MDM2/MDMX (Phe3, Trp7) were predicted to be buried within the protein (Fig. 4A, second and fourth panels).

We then employed molecular dynamics simulations using predicted structures by AlphaFold 3 to evaluate the impact of the truncation of the C-terminal amino acids in LOV2 on the dynamics of the PMI/PMI-M3 peptides (Fig. 4B and Movie 5). The simulation results indicated that the fusion to the Jα helix-truncated LOV2 restricted the movement of the PMI/PMI-M3 peptides and maintained tightly packed conformation, compared to fusion with the full-length LOV2. To quantify the flexibility of the PMI/PMI-M3 peptide, we calculated the average of the root mean square fluctuation (RMSF) value for all residues in the PMI/PMI-M3 peptide, exhibiting reduced RMSF values for the Jα helix-truncated LOV2 fusion than those for full-length LOV2 (Fig. 4C; RMSF values for all residues in the LOV2-PMI module are shown in Fig. S4A). Next, we evaluated the exposure of residues critical for MDM2/MDMX interaction to the protein surface using solvent accessible surface area (SASA) (Fig. 4D; SASA values for all residues in PMI/PMI-M3 are shown in Fig. S4B). The SASA value of N-terminal Phe3 decreased in both PMI and PMI-M3 when fused to the Jα helix-truncated LOV2 (Fig. 4D, left). The SASA value of Trp7 decreased only when the Jα helix-truncated LOV2 was fused to PMI-M3 (Fig. 4D, middle), and no clear difference was observed for C-terminal Leu10 (Fig. 4D, right). Taken together, these results strongly suggested that fusing to Jα helix-truncated LOV2 restricts the overall flexibility of the PMI/PMI-M3 peptides. Furthermore, the exposure of residues important for interaction with MDM2/MDMX to the protein surface is reduced. These changes likely prevent the LOV2-PMI modules from binding to MDM2/MDMX under dark conditions.

### Further screening of LOV2-PMI modules with different PMI mutants

We discovered that fusing PMI peptides with LOV2 truncated by 6 or 7 residues in the Jα helix yielded LOV2-PMI modules with remarkable light responsivity. Next, we aimed to further improve this responsiveness by testing a series of mutant peptides previously developed during the optimization of PMI/PMI-M3 peptides (Li et al., 2010, 2021). We selected several PMI variants reported to possess distinct *in vitro* affinities for MDM2/MDMX (Fig. 5A). Additionally, we tested peptide sequences derived from the MDM2/MDMX binding site of p53 (Schon et al., 2002). These peptide sequences share three residues (Phe3, Trp7, and Leu10) critical for the MDM2/MDMX interaction. We constructed an expression library by fusing these peptides to LOV2 with 6- or 7-residue truncation in the Jα helix (Fig. 5B). Using this library, we identified several LOV2-PMI fragments showing marked differences in binding activity toward MDM2/MDMX between the *dsm* and *lsm* (Fig. 5C, D; full screening data are shown in Fig. S5A, B). Notably, the LOV2(–6Jα)-PMI(E5A) mutant showed a difference in binding activity between *dsm* and *lsm* comparable to that of LOV2(−7Jα)-PMI-M3, despite the E5A peptide itself having lower affinity for MDM2/MDMX. We also observed that binding activities differed by up to several-fold between *dsm* and *lsm* even when using sequences derived from the p53 as MDM2/MDMX binding peptides (Fig. 5D and Fig. S5B).

**Figure 5.**
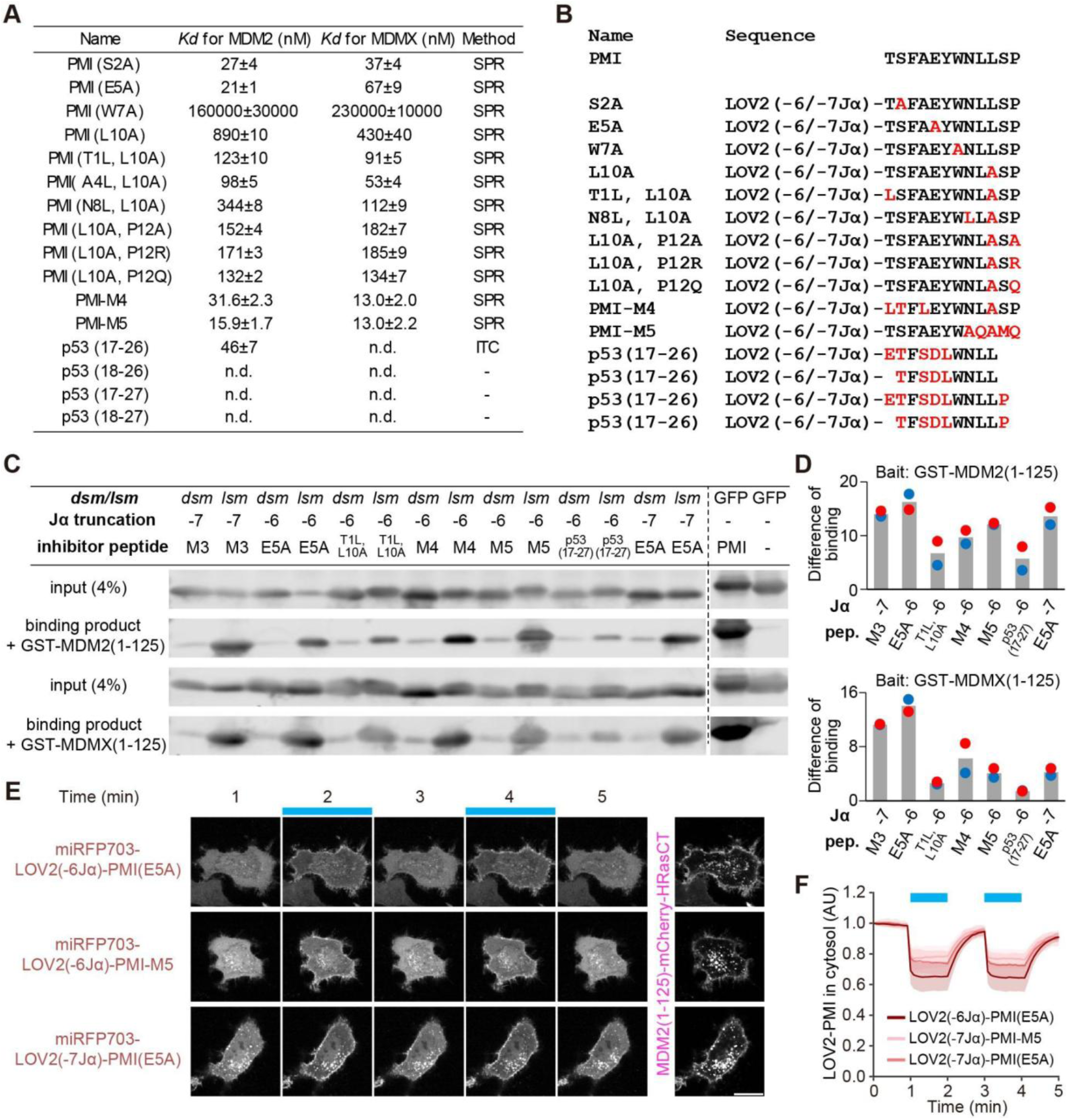
Expanded screening of LOV2-PMI actuator using an inhibitory peptide library. A. Table of inhibitory peptide variants with different *in vitro* affinities for MDM2/MDMX. SPR, surface plasmon resonance; ITC, isothermal titration calorimetry. B. Expression library of the LOV2-inhibitory peptide variants. Residues differing from the original PMI sequence are shown in red. C. Representative western blot images for the detection of each LOV2-inhibitory peptide fragments bound to GST–MDM2(1–125) or GST-MDMX(1–125). GFP-PMI and GFP serve as positive and negative control prey fragments, respectively. D. Comparison of binding activities between the *dsm* and *lsm* of each LOV2-inhibitory peptide fragment toward GST–MDM2(1–125) (upper) and GST-MDMX(1–125) (lower). Blue and red points represent experimental replicates. E. Light-dependent translocation of the Opto-MDMi (LOV2-PMI) actuator fragment obtained by the *in vitro* binding assay. Blue boxes indicate periods of light illumination. Scale bar, 20 µm. F. Quantification of the Opto-MDMi (LOV2-PMI) actuator levels in the cytoplasm. Plots show the mean ± s.d. (LOV2(–6Jα)-PMI(E5A), n = 40 cells; LOV2(–7Jα)-PMI-M5, n = 20 cells; LOV2(–7Jα)-PMI(E5A), n = 30 cells).

We next verified whether the LOV2(−6Jα)-PMI(E5A), LOV2(−6Jα)-PMI-M5, and LOV2(−7Jα)-PMI(E5A) fragments, which showed good performance in the *in vitro* binding assay, interacted with MDM2 in a light-dependent manner within living cells. As expected, all three LOV2-PMI fragments clearly translocated to the plasma membrane upon blue light illumination (Fig. 5E, F and Movie 6). No substantial differences in the translocation dynamics were observed among these variants. Consistent with the *in vitro* results, LOV2(–6Jα)-PMI(E5A) exhibited the highest efficiency for plasma membrane localization. Therefore, we selected the LOV2(–6Jα)-PMI(E5A) module for subsequent experiments.

### Light-dependent activation of p53 by Opto-MDMi (LOV2-PMI)

Next, we investigated whether the obtained LOV2-PMI chimera modules could actually activate endogenous p53. For this purpose, we established cell lines stably expressing fluorescently tagged LOV2-PMI fragments (Fig. 6A and Fig. S6A). In this system, miRFP703-NLS-LOV2-PMI actuator fragments are expressed in the nucleus under dark conditions (Fig. 6A, left). Upon blue light illumination, the actuator fragments expose the PMI/PMI-M3 peptides, which are expected to induce p53 activation by inhibiting MDM2/MDMX (Fig. 6A, right). This optogenetic system is hereafter referred to as Opto-MDMi (LOV2-PMI). Transcriptional activation of p53 was induced by all tested LOV2-PMI modules upon blue light illumination (Fig. 6B, C and Movie 7). Of note, the increase in p53 reporter activity observed under dark conditions in the Opto-MDMi (LOVTRAP) system was hardly observed in the Opto-MDMi (LOV2-PMI) system (compare Fig. 1G and Fig. 6C). Furthermore, quantification of actuator expression in cells before cell sorting revealed an increase in the population of cells expressing higher levels of the actuators, likely due to reduced interaction between the actuator and MDM2/MDMX under basal conditions (compare Fig. S1F and Fig. S6B). The transcription reporter of p53 increased approximately 3.1-, 3.7-, and 2.8-fold for LOV2(–6Jα)-PMI, LOV2(–7Jα)-PMI-M3, and LOV2(−6Jα)-PMI(E5A), respectively, outperforming the results obtained with the Opto-MDMi (LOVTRAP) system (Fig. 6D, see also Fig. 1H). We also quantified the temporal changes in the expression levels of the actuator fragments (Fig. S6C). The expression of the negative control fragment slightly increased. In contrast, all actuator fragments showed a tendency for their expression levels to gradually decrease upon blue light illumination, suggesting that the light-dependent interaction with MDM2 induces the degradation of the actuator fragments themselves. Interestingly, the fragment containing PMI (E5A), which exhibited the lowest *in vitro* binding affinity to MDM2/MDMX among these three, showed the largest decrease for unknown reasons. Nevertheless, our results clearly demonstrate that the Opto-MDMi (LOV2-PMI) actuator possesses the ability to activate the p53 pathway.

**Figure 6.**
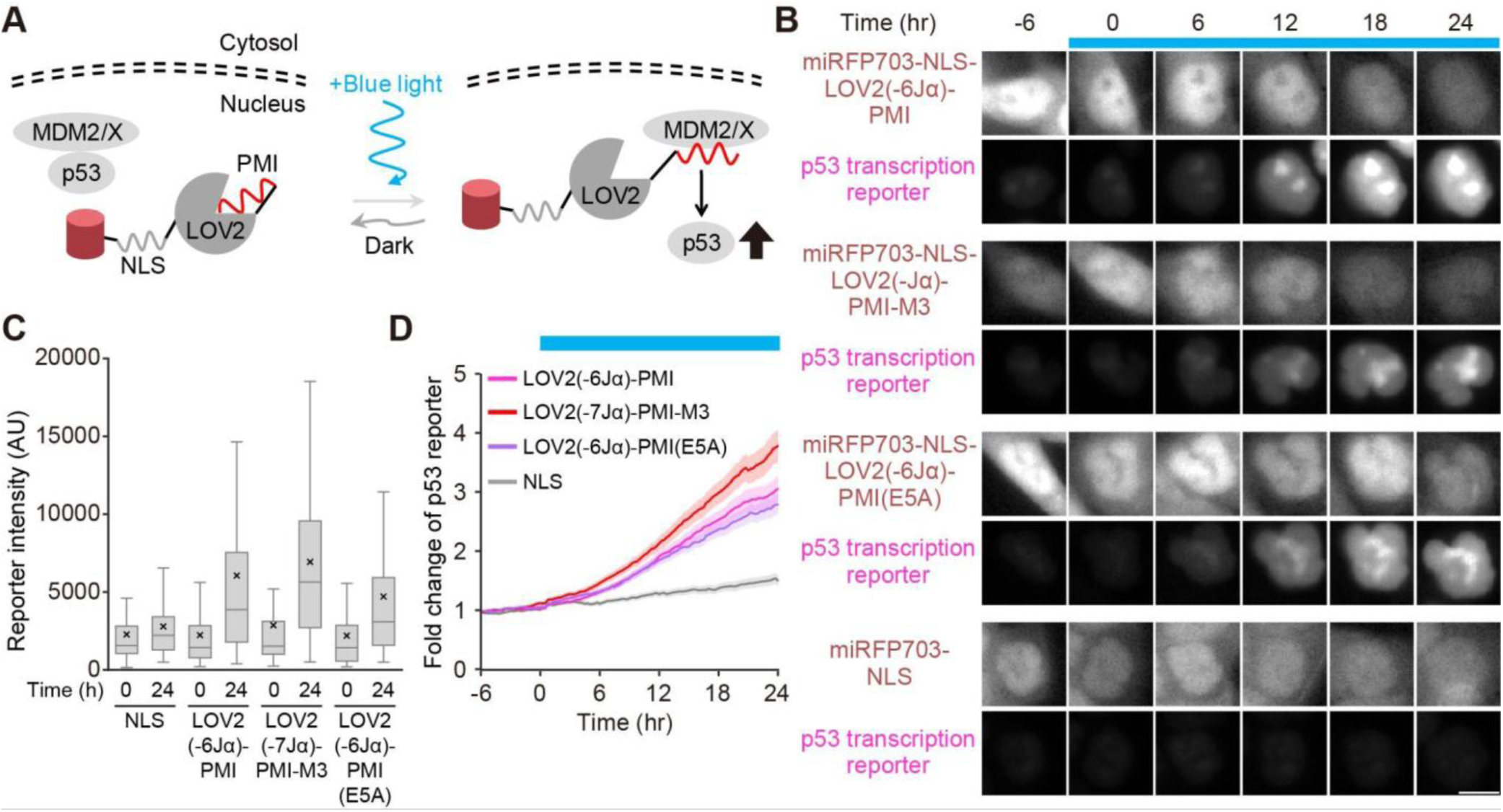
Light-dependent transcriptional activation of p53 by Opto-MDMi (LOV2-PMI) A. Schematic illustration of the Opto-MDMi (LOV2-PMI) system expressed in cultured cells. B. Light-dependent changes in the actuator and p53 transcription reporter signals. The blue boxes indicate the time points at which light illumination was applied. Scale bar, 10 µm. C. Distribution of the fluorescence intensity of the p53 transcription reporter at 0 hours and 24 hours from the start of blue light illumination. Horizontal lines and crosses indicate the medians and means of the distribution, respectively. Horizontal lines and crosses indicate the medians and means, respectively. Boxes represent the 25th and 75th percentiles; whiskers indicate the range between the maximum and minimum values excluding outliers. D. Temporal changes in the p53 transcription reporter in each cell line. The plot shows the mean ± s.e.m. (LOV2(–6Jα)-PMI, n = 139 cells; LOV2(–7Jα)-PMI-M3, n = 116 cells; LOV2(–6Jα)-PMI(E5A), n = 170 cells; NLS, n = 82 cells).

### Combination of Opto-MDMi (LOVTRAP) and Opto-MDMi (LOV2-PMI) system

Finally, we examined whether combining Opto-MDMi (LOVTRAP) and Opto-MDMi (LOV2-PMI) allows for more efficient activation of p53. We designed a system capable of dual control by replacing the actuator fragment in Opto-MDMi (LOVTRAP) with the Opto-MDMi (LOV2-PMI) actuator (Fig. 7A). In this system, simply called Opto-MDMi, the actuator fragment binds to the localizer, and the activity of the PMI peptide is restricted by steric hindrance from LOV2 under dark conditions (Fig. 7A, left). Upon blue light illumination, the actuator fragment exposes the PMI peptide and simultaneously dissociates from the localizer (Fig. 7A, right). The released actuator fragment then translocates to the nucleus, where it is expected to induce p53 activation. A potential obstacle for the Opto-MDMi system is that both the Opto-MDMi (LOVTRAP) localizer and the Opto-MDMi (LOV2-PMI) actuator contain LOV2 domains. Therefore, the Zdk1 fragment might also interact with the LOV2 domain within the Opto-MDMi (LOV2-PMI) actuator. Previous reports have shown that the C-terminal sequence of LOV2 is crucial for its interaction with Zdk1, and modifying this sequence results in the loss of Zdk1-binding ability (Wang et al., 2016). To confirm whether the Opto-MDMi (LOV2-PMI) actuators interact with Zdk1, we designed a new experimental system (Fig. 7A). In this system, the LOV2-PMI modules were anchored to mitochondria instead of the LOVTRAP localizer, and a Zdk1-NLS fragment was expressed in the cell. This allows us to determine the presence of interaction based on the localization of Zdk1-NLS. We expressed five variants (LOV2(–6Jα)-PMI, LOV2(–7Jα)-PMI-M3, LOV2(–6Jα)-PMI(E5A), LOV2(–6Jα)-PMI-M5, and LOV2(–7Jα)-PMI(E5A)) and tested their localization (Fig. S7B). As a result, interaction with Zdk1 was completely abolished in all variants except LOV2(–7Jα)-PMI(E5A) (Fig. S7C). Only LOV2(–7Jα)-PMI(E5A) showed a clear interaction with Zdk1; however, unlike wild-type LOV2, this interaction was not released by light illumination. These results showed that LOV2-PMI actuators, except for LOV2(–7Jα)-PMI(E5A), could be successfully combined with the protein localization manipulation via LOVTRAP.

**Figure 7.**
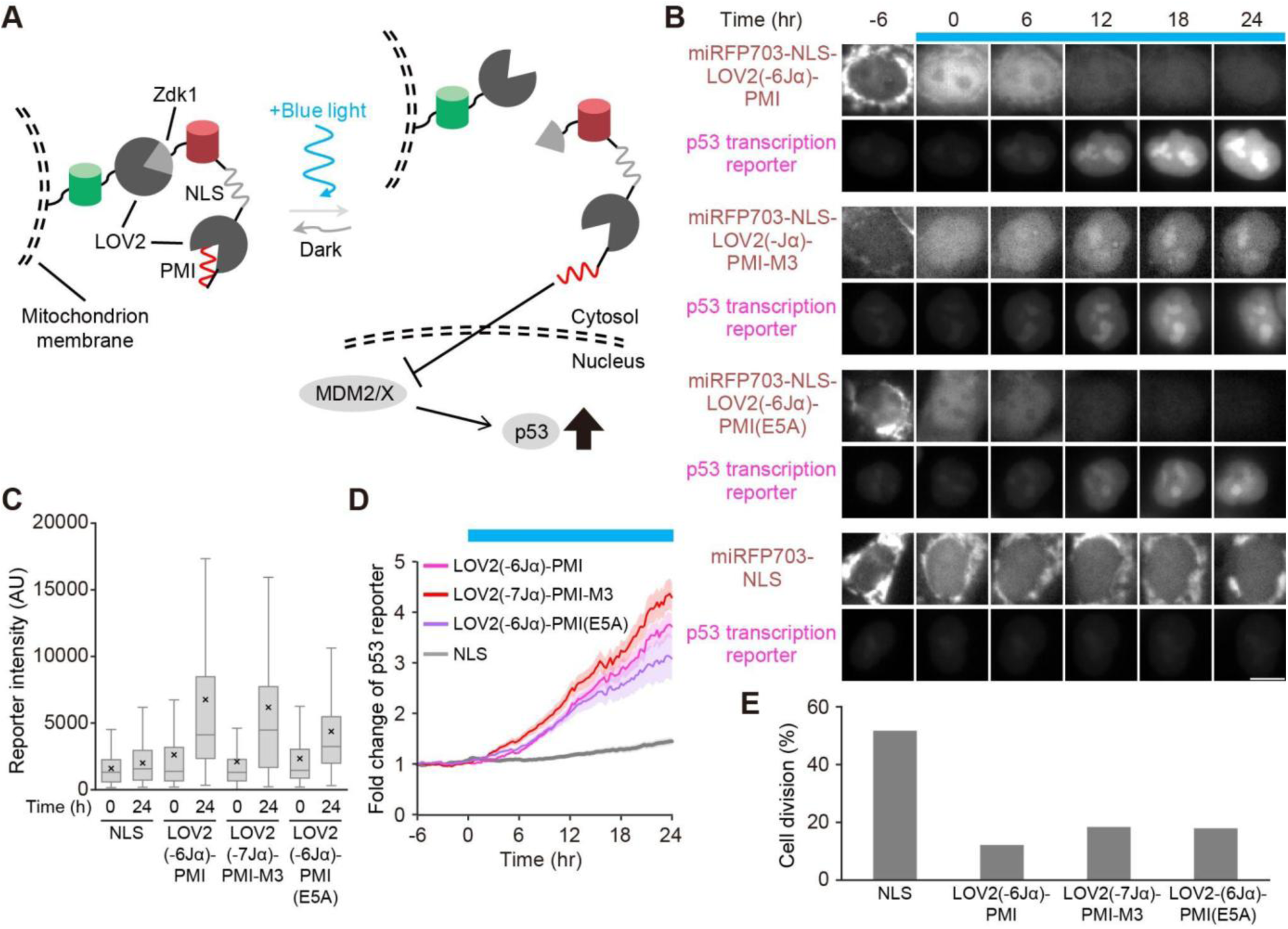
Light-dependent transcriptional activation of p53 by combined Opto-MDMi system. A. Schematic illustration of the combination of Opto-MDMi (LOVTRAP) and Opto-MDMi (LOV2-PMI). B. Light-dependent changes in Opto-MDMi actuator localization and p53 transcription reporter signals. The blue boxes indicate the time points at which light illumination was applied. Scale bar, 10 μm. C. Distribution of the fluorescence intensity of the p53 transcription reporter at 0 hours and 24 hours from the start of blue light illumination. Horizontal lines and crosses indicate the medians and means of the distribution, respectively. Horizontal lines and crosses indicate the medians and means, respectively. Boxes represent the 25th and 75th percentiles; whiskers indicate the range between the maximum and minimum values excluding outliers. D. Temporal changes in the p53 transcription reporter in each cell line. The plot shows the mean ± s.e.m. (LOV2(–6Jα)-PMI, n = 181 cells; LOV2(–7Jα)-PMI-M3, n = 185 cells; LOV2(–6Jα)-PMI(E5A), n = 123 cells; NLS, n=209 cells). E. Quantification of cell division events during the period from 0 to 24 hours in panel D.

We established stable cell lines co-expressing the Opto-MDMi localizer, actuator, and p53 reporter. We quantified actuator expression in these cell lines before cell sorting and confirmed that the expression levels were comparable to the cell line expressing only Opto-MDMi (LOV2-PMI) actuators (compare Fig. S6B and Fig. S7D). We sorted cell populations showing high expression of each fragment and used them in the following experiments. Upon blue light illumination, transcriptional activation of p53 was observed in these cells (Fig. 7B, C and Movie 8). The transcription reporter of p53 increased approximately 3.7-, 4.3-, and 3.1-fold for LOV2(–6Jα)-PMI, LOV2(–7Jα)-PMI-M3, and LOV2(–6Jα)-PMI(E5A), respectively (Fig. 7D). Compared to Opto-MDMi (LOV2-PMI) alone, this represented increases of 22%, 13%, and 10% (Fig. 6D and Fig. 7D). Although the significance of these changes remains open to discussion due to intercellular variation, imaging noise, and other factors, the consistent observation of further increases in reporter intensity across all three cell lines suggests improved function resulting from the combination of the two optogenetic systems. In contrast, these cell lines showed minimal p53 transcriptional activation in the dark (Fig. S7E). This may be due to cells spontaneously activating p53 by the expression of the PMI peptide. We also quantified the cell division events during 24-hour light illumination period and found that cell proliferation was suppressed in all Opto-MDMi expressing cells compared to the negative control (Fig. 7E). This growth inhibition was not observed at all under dark conditions, demonstrating that the Opto-MDMi system efficiently restricts cell cycle progression through light-dependent p53 activation (Fig. S7F). Overall, our findings demonstrate that the Opto-MDMi enables the activation of endogenous p53 to a degree capable of altering cellular phenotypes.

## Discussion

In this study, we successfully established Opto-MDMi, an optogenetic platform that controls the activity of endogenous p53 via light. Opto-MDMi enables light-dependent activation of p53 by simultaneously regulating the subcellular localization and the inhibitory activity of peptides that inhibit the interaction between p53 and MDM2/MDMX. This multi-layer control architecture, achieved through the integration of two or more optogenetic modules, may serve as a general principle for designing optogenetic tools with an improved dynamic range when the performance of individual modules is limited. Indeed, several studies have shown that combining a light-induced nuclear translocation system with cytoplasmic anchoring reduces leaky nuclear localization in the dark, achieving tight light-dependent nuclear translocation (Redchuk et al., 2017; Yumerefendi et al., 2018; Chen et al., 2020). In our study, combining Opto-MDMi (LOVTRAP) with Opto-MDMi (LOV2-PMI) yielded the gain of p53 activation (Fig. 7). Given that the interaction between the LOV2 core region and the Jα helix is essentially in equilibrium—with approximately 1.6% remaining undocked even under dark conditions—multi-layer control appears to be an effective strategy for improving the performance of optogenetic tools by suppressing undesired basal leakage (Yao et al., 2008).

The most notable feature of the Opto-MDMi system is its ability to manipulate the activity of endogenous p53. Hub proteins in signaling pathways, such as p53, often possess multiple functions. Therefore, constitutively expressing domains derived from such proteins may cause unexpected non-physiological effects, potentially limiting their application. In fact, p53 is known to have transcription-independent functions that induce apoptosis through the interaction with anti-apoptotic proteins (Caelles et al., 1994; Wagner et al., 1994; Haupt et al., 1995; Marchenko et al., 2000; Mihara et al., 2003). This is consistent with the observation that a certain fraction of cell death occurs even under dark conditions in Opto-p53, an optogenetics system that exogenously expresses p53 fragments (Tsuruoka et al., 2025). In contrast, optogenetic systems based on short functional peptides, such as the one developed in this study, do not require the expression of the target signaling factor fragment itself. This approach is expected to minimize side effects while precisely perturbing the desired target.

Photocaging short functional peptides onto LOV2 is a promising approach for generating compact light-responsive actuators. Consequently, chimeric modules combining LOV2 with various functional peptides have been developed, though general design principles have not yet been established. In past successful attempts, optogenetic systems controlling nuclear import/export or protein-protein interactions were developed by incorporating peptides into the Jα helix, leveraging sequence homology (Lungu et al., 2012; Yumerefendi et al., 2015, 2016; Niopek et al., 2014, 2016). However, this strategy requires the presence of peptides with substantial sequence homology to the Jα helix. Alternatively, reports using JNK inhibitory peptides have employed a strategy of truncating the inhibitory peptide itself (Melero-Fernandez de Mera et al., 2017). However, truncating residues on the peptide side is likely to reduce its intrinsic activity, making it difficult to regard as a generally applicable approach (Fig. S2A-D). In contrast, the strategy demonstrated in the Opto-MDMi (LOV2-PMI) system—truncating the Jα helix of LOV2—preserves the fully functional peptide (Fig. 2). This makes it a preferable approach for introducing light responsivity to functional peptides. The binding between the LOV2 core domain and the Jα helix under dark conditions is largely mediated by hydrophobic interactions at their interface (Harper et al., 2004). Among these, Ile539 in the Jα helix makes the largest contribution, suggesting that the Jα helix is pinned to the core domain by this residue. Therefore, fusing a peptide to a truncated LOV2 likely restrains the movement of peptides as well (Fig. 4). Thus, a general strategy for photocaging functional peptides within LOV2 could be to position the critical residues of the peptide toward the LOV2 core domain (thereby hiding them) while deleting as many residues as possible from the Jα helix side.

The design of the optogenetic tool presented in this study has the potential to be extended to the spatiotemporal control of various other ubiquitin ligase activities by modifying the peptide sequence that binds to the substrate recognition domain. However, we observed a gradual decrease in actuator expression levels during light illumination in the Opto-MDMi (LOV2-PMI) system, suggesting that degradation may have occurred once the actuator interacts with MDM2 (Fig. S6C). Therefore, to apply our approach to more robustly manipulate other ubiquitin ligases, it will be necessary to engineer a degradation-resistant LOV2, for example, by mutating lysine residues exposed on the protein surface. Ultimately, combining such systems with patterned optogenetic stimulation devices to enable subcellular-scale manipulation of ubiquitin ligase activity will lead to a deeper understanding of the biological processes involving them. Thus, the Opto-MDMi system serves as a framework for the future development of the tools to manipulate ubiquitin ligase activity.

## Materials and Methods

### Construction of plasmids

All plasmids used in this study (except for the expression library used in the *in vitro* binding assay) are summarized in Table S1, along with Benchling links to the plasmid sequences and maps. The oligonucleotides for PCR and DNA sequencing were obtained from FASMAC. For the p53 transcription reporter vector, we used a vector previously developed in our laboratory (Tsuruoka et al., 2025). The cDNAs of the p53 binding domains of human MDM2 and MDMX were cloned by PCR using total cDNA derived from HCT116 cells as a template. The AsLOV2(404-546) and Zdk1 genes were obtained from pTriEx-NTOM20-LOV2 (plasmid#81009: Addgene) and pTriEx-mCherry-Zdk1-VAV2 DH/PH/C1 (plasmid#81060: Addgene), respectively (Wang et al., 2016). The cDNA of HRasCT was obtained from pCAGGS-hPhyB-mCherry-HRasCT (plasmid#100281: Addgene) (Uda et al., 2017). Short functional sequences such as NLS and PMI peptides were inserted using DNA fragments generated by oligo DNA annealing. The expression vectors for the PMI-truncated LOV2-PMI library (Table S2) were also constructed via oligo DNA annealing method. The expression vectors for the Jα helix-truncated LOV2-PMI library (Table S3), the PMI-truncated LOV2(V416L)-PMI expression library (Table S4), and the LOV2-PMI mutant expression library (Table S5) were constructed by site-directed mutagenesis using inverse PCR. All other expression vectors were constructed by restriction enzyme digestion and ligation, or by Gibson Assembly using NEBuilder HiFi DNA Assembly (New England Biolabs).

### Cell culture

HeLa and HCT116 cells were obtained from the American Type Culture Collection (ATCC; Rockville, MD, USA). HeLa cells were cultured in DMEM (high glucose) (08459-64: Nacalai) supplemented with 10% fetal bovine serum (FBS; 175012: NICHIREI). HCT116 cells were cultured in RPMI 1640 Medium (ATCC modification) (A10491-01: Thermo Fisher Scientific) supplemented with 10% FBS. All cells were maintained in a humidified atmosphere of 5% CO_2_ at 37°C.

### Transfection

All transient expression experiments were performed using HeLa cells. Cells were seeded on a collagen-coated 4-well glass-bottom dish (The Greiner Bio-One) one day prior to transfection, and were transfected with total 1-2 μg of plasmids by using Polyethyleneimine ‘Max’ MW 40,000 (Polyscience Inc., Warrington, PA, USA). In Figure 1D and S1B, expression vectors for Opto-MDMi (LOVTRAP) localizer, Opto-MDMi (LOVTRAP) actuator, and H1-mCherry (nuclear marker) were transfected at a 2:1:1 ratio. In Figure 3B and 5E, MDM2(1-125) fragment and Opto-MDMi (LOV2-PMI) actuator were transfected at a 1:2 ratio. In Figure S7C, LOV2-PMI fragment and Zdk1 fragment were transfected at a 2:1 ratio. Time-lapse imaging was started 1 day after transfection.

### Establishment of stable cell lines

All stable cell line experiments were performed using HCT116 cells. Cells were transfected with PiggyBac donor vectors and PiggyBac transposase-expressing vectors at a ratio of 3:1 (Yusa et al., 2009, 2011). Nucleofector IIb (Lonza, Basel) electroporation system was used for transfection according to the manufacturers’ instructions (D-032 program) with a house-made DNA- and cell-suspension solution (4 mM KCl, 10 mM MgCl_2_, 107 mM Na_2_HPO_4_, 13 mM NaH_2_PO_4_, 11 mM HEPES pH 7.75) (Yamamoto et al., 2021). One day after transfections, cells were treated with 1 μg/mL puromycin (InvivoGen, San Diego, CA), 10 μg/mL blasticidin S (InvivoGen), or 1000 μg/mL G418 (InvivoGen) for drug selection. Established cell lines were sorted by MA900 Multi-Application Cell Sorter (SONY) or BD FACSAriaII Flow Cytometer (BD) based on the fluorescence signal from localizer and actuator fragments. Time-lapse imaging for stable cell lines was started 2 days after seeding on a collagen-coated 4-well glass-bottom dish. For Fig. S1D and E, the stable cell line harboring p53 transcription reporter was seeded into a 96-well flat-bottom microplate (The Greiner Bio-One).

### Live-cell fluorescence imaging

Live-cell imaging was performed using epifluorescence inverted microscopes (IX83; Olympus, Tokyo). For spinning-disk confocal microscopy (Fig. 1E, 3B, 5E, and S1B), the microscope was equipped with a spinning-disk confocal unit (CSU-W1; Yokogawa Electric Corporation), an sCMOS camera (ORCA Fusion BT; Hamamatsu Photonics), and an oil-immersion objective lens (UPLXAPO 60X, NA = 1.42, WD = 0.15 mm; Olympus). The excitation lasers and emission filters used were as follows: Excitation laser, 488 nm for mNeonGreen, 561 nm for mCherry, and 640 nm for miRFP703; excitation dichroic mirror, DM405/488/561/640 for mNeonGreen, mCherry, and miRFP703; emission filters, 525/50 for mNeonGreen, 617/73 for mCherry, and 685/40 for miRFP703 (Yokogawa Electric).

For wide-field fluorescence microscopy (Fig. 1G, 6B, and 7B), the microscope was equipped with a Prime sCMOS camera (Photometrics) and dry objective lens (UPLXAPO 40X). A Spectra X light engine (Lumencor) was used as the illumination light source with the following setting: Excitation wavelength, 475/28 for mNeonGreen, 580/20 for mScarlet-I, and 632/22 for miRFP703; dichroic mirror, FF409/493/573/652/759-Di01-25.8×37.8 for all fluorescent proteins; emission filters, 520/28 for mNeonGreen, 641/75 for mScarlet-I, and 664/long pass for miRFP703 (Semrock). For optogenetic stimulation, blue LED light (450 nm) (LED-41VIS450; OptoCode Corp., Japan) was continuously delivered from the top of the stage.

For the experiments shown in Fig. S1D and E, an ImageXpress Micro XLS (Molecular Devices) was employed, equipped with an sCMOS camera (Zyla 5.5; Andor) and dry objective lens (Plan Fluor 10X, NA = 0.30, WD = 16 mm; Nikon). The illumination light source was a SOLA SE2 (Lumencor). The illumination settings were as follows: Excitation wavelength, 562/40 for mScarlet-I; excitation dichroic mirror, 350-585 (R)/601-950 (T) for mScarlet-I; emission filters, 624/40 for mScarlet-I (Semrock).

### Image Analysis

All imaging data were analyzed using Fiji/ImageJ (Schindelin et al., 2012). Background subtraction was performed on all images using the rolling-ball method, followed by the generation of stacked images. Nuclear regions were detected and cell trajectories were tracked using LIMTracker, a Fiji tracking plugin (Aragaki et al., 2022), which allowed for the quantification of temporal changes in fluorescence profiles for individual cell traces. Nuclear regions were segmented based on either the H1-mCherry signal (Fig. 1F) or the p53 transcriptional reporter signal (mScarlet-I-3xNLS; Fig. 1G, 1H, 6C, 6D, 7C, 7D). For the data of Fig. 3C, 5F, S1C, and S3C, regions of interest (ROIs) were manually defined, and fluorescence intensity changes within the ROI were quantified. For the datasets in Fig. S1D and S1E, the Fiji plugin StarDist (Schmidt et al., 2018) was employed for automated nuclei detection at each time point, followed by quantification of the fluorescent signal in the nuclei. All the quantified data were analyzed and visualized using Python with the seaborn and matplotlib libraries or Microsoft Excel software.

### Expression and purification of recombinant proteins

Bait protein fragments (GST, GST-MDM2(1-125), and GST-MDMX(1-125)) were expressed in *E. coli* BL21(DE3) cells using the pCold vector system (Qing et al., 2004). Competent *E. coli* BL21(DE3) cells were transformed with pCold expression vectors containing the desired gene cassettes. Grown colonies were picked and inoculated into 2 mL of LB medium supplemented with 0.1 mg/mL ampicillin and cultured overnight at 37°C. The next day, the culture was collected, mixed with a final concentration of 7% DMSO, and stored at −80°C as a frozen stock.

For protein expression, this stock was re-inoculated into 1 mL of LB medium (0.1 mg/mL ampicillin) and cultured at 37°C. After several hours, the culture was transferred to 100 mL of LB medium (0.1 mg/mL ampicillin) and cultured until the optical density (OD) reached 0.4–0.8. The culture was then rapidly cooled in ice water, shaken at 15°C for 30 minutes, and induced with a final concentration of 0.3 mM IPTG overnight at 15°C. The bacterial cells were harvested by centrifugation, resuspended in 10 mL of purification buffer (1 mM dithiothreitol, 300 mM NaCl, 0.1% Triton X-100 in PBS), and lysed using an ultrasonic homogenizer VP-300N (TAITEC). The lysate was centrifuged, and the resulting supernatant was incubated overnight at 4°C with 1 mL of Glutathione-Sepharose 4B beads (Cytiva) pre-equilibrated with purification buffer. After incubation, the supernatant was removed, and the beads were washed three times with purification buffer. Finally, an equal volume of purification buffer was added to the beads. To verify successful expression and purification, 10 μL of the bead suspension was mixed with 40 μL of 1 × SDS sample buffer (62.5 mM Tris-HCl pH 6.8, 11.9% glycerol, 2% SDS, 5% dithiothreitol) and boiled at 95°C for 5 minutes. Subsequently, 10 μL of the boiled samples were analyzed by SDS-polyacrylamide gel electrophoresis (SDS-PAGE) followed by Coomassie Brilliant Blue (CBB) staining (Fig. 2C). The CBB-stained gel was imaged using an Odyssey Infrared Imaging System (LI-COR). The purified bait protein fragments immobilized on the beads were used for subsequent *in vitro* binding assays.

### *in vitro* binding assay

For the expression of prey protein fragments (LOV2-PMI library), *E. coli* BL21(DE3) cells were transformed with pCold expression vectors following the same procedure as for the bait protein, and frozen stocks were prepared. These stocks were re-inoculated in 2 mL of LB medium (0.1 mg/mL ampicillin) and cultured at 37°C until the OD reached 0.4-0.8. The cultures were then rapidly cooled in ice water, shaken at 15°C for 30 minutes, induced with a final concentration of 0.3 mM IPTG, and incubated overnight at 15°C. Bacterial cells were harvested by centrifugation and lysed using freeze-thaw method. Specifically, cell pellets were resuspended in 250 µL/samples of resuspension buffer (50 mM Tris-HCl pH 7.5, 500 mM NaCl, 1 mM dithiothreitol), frozen at −20°C for 30 minutes, and thawed at 4°C for 2 hours. Subsequently, 125 µL/sample of lysis buffer (50 mM Tris-HCl pH 7.5, 500 mM NaCl, 1 mM dithiothreitol, 25 µg/sample lysozyme (Thermo), 31.25 Unit/sample benzonase nuclease (Merck), 0.5% Triton X-100) was added to the cell suspension, followed by incubation at 37°C for 1 hour. After incubation, the samples were centrifuged, and the supernatant was collected as the cell lysate containing the prey protein fragments. For analysis, 15 μL of these cell lysates were mixed with 15 μL of 2 × SDS sample buffer (130 mM Tris-HCl, pH 6.8, 23.8% glycerol, 4% SDS, 10% dithiothreitol) and boiled at 95°C for 5 minutes to serve as the “input” fraction. The remaining lysates were incubated overnight at 4°C with 20 µL/sample of the bait protein fragment immobilized on Glutathione Sepharose 4B beads. The following day, the beads were washed three times with resuspension buffer. Finally, the beads were mixed with 1 × SDS sample buffer and boiled at 95°C for 5 minutes to serve as the “binding product” fraction.

### Western blotting

The prepared “input” and “binding product” samples were separated by SDS-PAGE, and transferred onto polyvinylidene difluoride (PVDF) membranes. Proteins were detected by using an Odyssey Infrared Imaging System (LI-COR). The primary antibody used was a rabbit monoclonal anti-T7-tag (D9E1X; 1:1000, Cell Signaling Technology, #13246). The secondary antibody was an IRDye 680RD goat anti-rabbit antibody (1:5000, LI-COR, 926-68071). Blotting images were quantified by using Image Studio software (LI-COR).

### Molecular dynamics simulation

The system was composed of a LOV2-PMI fragment (LOV2-PMI, LOV2(−6Jα)-PMI, LOV2-PMI-M3, or LOV2(−7Jα)-PMI-M3) predicted by AlphaFold 3 (Abramson et al., 2024) immersed in 150 mM KCl solution (see Table S6 for molecular composition). The AMBER ff19SB (Tian et al., 2020), OPC (Izadi et al., 2014), and Li and Merz (Sengupta et al., 2021) force fields were employed for the protein, water molecules, and ions, respectively. All histidine residues were protonated at the epsilon-carbon.

After 1,000 step energy minimization (where necessary), molecular dynamics (MD) simulations were performed using AMBER24 (Amber). First, a 40-ps MD simulation was conducted under constant temperature (300 K) and pressure (1 bar) condition. Subsequently, a 1-ns MD simulation with constant temperature (300 K) and volume condition was performed to equilibrate the system. The Berendsen thermostat and barostat (Berendsen et al., 1984) were used to control the temperature and pressure, respectively. The length of bonds having hydrogen atoms was kept constant using the SHAKE algorithm (Ryckaert et al., 1977), enabling the use of time step of 2 fs. The periodic boundary condition was applied, and long-range electrostatic interactions were calculated by the particle mesh Ewald method (Essmann et al., 1995) with a 10 Å cutoff in real space. For the production runs, MD simulations were carried out for 1 μs under constant temperature (300 K) and volume condition. The results of the MD simulation were visualized using VMD (Humphrey et al., 1996). Root-mean-square fluctuation (RMSF) values for each amino acid residue were calculated using in-house code based on the MD simulation results. Solvent-accessible surface area (SASA) was calculated using the following site (http://cib.cf.ocha.ac.jp/bitool/ASA/). More specifically, SASA was calculated from the structural data extracted every 100 frames, and then the average values were calculated for each MD trajectory.

## Acknowledgements

We thank all members of the Aoki Laboratory for their helpful discussions and assistance. We are grateful to E. Ebine-Sato, K. Onoda, M. Hirao, Y. Tomizawa, K. Takakura and T. Uesugi for their technical support. The MD simulations were carried out on the supercomputers at the Research Center for Computational Science in Okazaki, Japan (Project: 25-IMS-097).

## References

Abramson, J., J. Adler, J. Dunger, R. Evans, T. Green, A. Pritzel, O. Ronneberger, L. Willmore, A.J. Ballard, J. Bambrick, S.W. Bodenstein, D.A. Evans, C.-C. Hung, M. O’Neill, D. Reiman, K. Tunyasuvunakool, Z. Wu, A. Žemgulytė, E. Arvaniti, C. Beattie, O. Bertolli, A. Bridgland, A. Cherepanov, M. Congreve, A.I. Cowen-Rivers, A. Cowie, M. Figurnov, F.B. Fuchs, H. Gladman, R. Jain, Y.A. Khan, C.M.R. Low, K. Perlin, A. Potapenko, P. Savy, S. Singh, A. Stecula, A. Thillaisundaram, C. Tong, S. Yakneen, E.D. Zhong, M. Zielinski, A. Žídek, V. Bapst, P. Kohli, M. Jaderberg, D. Hassabis, and J.M. Jumper. 2024. Accurate structure prediction of biomolecular interactions with AlphaFold 3. Nature. 630:493–500.

Aragaki, H., K. Ogoh, Y. Kondo, and K. Aoki. 2022. LIM Tracker: a software package for cell tracking and analysis with advanced interactivity. Sci. Rep. 12:2702.

Baarlink, C., H. Wang, and R. Grosse. 2013. Nuclear actin network assembly by formins regulates the SRF coactivator MAL. Science. 340:864–867.

Berendsen, H.J.C., J.P.M. Postma, W.F. van Gunsteren, A. DiNola, and J.R. Haak. 1984. Molecular dynamics with coupling to an external bath. J. Chem. Phys. 81:3684–3690.

Caelles, C., A. Helmberg, and M. Karin. 1994. p53-dependent apoptosis in the absence of transcriptional activation of p53-target genes. Nature. 370:220–223.

Case, D.A., H.M. Aktulga, K. Belfon, I.Y. Ben-Shalom, J.T. Berryman, S.R. Brozell, D.S. Cerutti, T.E. Cheatham, III, G.A. Cisneros, V.W.D. Cruzeiro, T.A. Darden, N. Forouzesh, M. Ghazimirsaeed, G. Giambaşu, T. Giese, M.K. Gilson, H. Gohlke, A.W. Goetz, J. Harris, Z. Huang, S. Izadi, S.A. Izmailov, K. Kasavajhala, M.C. Kaymak, A. Kovalenko, T. Kurtzman, T.S. Lee, P. Li, Z. Li, C. Lin, J. Liu, T. Luchko, R. Luo, M. Machado, M. Manathunga, K.M. Merz, Y. Miao, O. Mikhailovskii, G. Monard, H. Nguyen, K.A. O’Hearn, A. Onufriev, F. Pan, S. Pantano, A. Rahnamoun, D.R. Roe, A. Roitberg, C. Sagui, S. Schott-Verdugo, A. Shajan, J. Shen, C.L. Simmerling, N.R. Skrynnikov, J. Smith, J. Swails, R.C. Walker, J. Wang, J. Wang, X. Wu, Y. Wu, Y. Xiong, Y. Xue, D.M. York, C. Zhao, Q. Zhu, and P.A. Kollman (2024), Amber 2024, University of California, San Francisco.

Chen, S.Y., L.C. Osimiri, M. Chevalier, L.J. Bugaj, T.H. Nguyen, R.A. Greenstein, A.H. Ng, J. Stewart-Ornstein, L.T. Neves, and H. El-Samad. 2020. Optogenetic Control Reveals Differential Promoter Interpretation of Transcription Factor Nuclear Translocation Dynamics. Cell Syst. 11:336–353.e24.

Dagliyan, O., M. Tarnawski, P.-H. Chu, D. Shirvanyants, I. Schlichting, N.V. Dokholyan, and K.M. Hahn. 2016. Engineering extrinsic disorder to control protein activity in living cells. Science. 354:1441–1444.

Essmann, U., L. Perera, M.L. Berkowitz, T. Darden, H. Lee, and L.G. Pedersen. 1995. A smooth particle mesh Ewald method. J. Chem. Phys. 103:8577–8593.

Guntas, G., R.A. Hallett, S.P. Zimmerman, T. Williams, H. Yumerefendi, J.E. Bear, and B. Kuhlman. 2015. Engineering an improved light-induced dimer (iLID) for controlling the localization and activity of signaling proteins. Proc. Natl. Acad. Sci. U. S. A. 112:112–117.

Halavaty, A.S., and K. Moffat. 2007. N- and C-terminal flanking regions modulate light-induced signal transduction in the LOV2 domain of the blue light sensor phototropin 1 from Avena sativa. Biochemistry. 46:14001–14009.

Harper, S.M., J.M. Christie, and K.H. Gardner. 2004. Disruption of the LOV-Jalpha helix interaction activates phototropin kinase activity. Biochemistry. 43:16184–16192.

Harper, S.M., L.C. Neil, and K.H. Gardner. 2003. Structural basis of a phototropin light switch. Science. 301:1541–1544.

Haupt, Y., R. Maya, A. Kazaz, and M. Oren. 1997. Mdm2 promotes the rapid degradation of p53. Nature. 387:296–299.

Haupt, Y., S. Rowan, E. Shaulian, K.H. Vousden, and M. Oren. 1995. Induction of apoptosis in HeLa cells by trans-activation-deficient p53. Genes Dev. 9:2170–2183.

He, L., Y. Zhang, G. Ma, P. Tan, Z. Li, S. Zang, X. Wu, J. Jing, S. Fang, L. Zhou, Y. Wang, Y. Huang, P.G. Hogan, G. Han, and Y. Zhou. 2015. Near-infrared photoactivatable control of Ca(2+) signaling and optogenetic immunomodulation. Elife. 4. doi:10.7554/eLife.10024.

Hoffmann, M.D., F. Bubeck, R. Eils, and D. Niopek. 2018. Controlling cells with light and LOV. Adv. Biosyst. 2:1800098.

Honda, R., H. Tanaka, and H. Yasuda. 1997. Oncoprotein MDM2 is a ubiquitin ligase E3 for tumor suppressor p53. FEBS Lett. 420:25–27.

Hongdusit, A., P.H. Zwart, B. Sankaran, and J.M. Fox. 2020. Minimally disruptive optical control of protein tyrosine phosphatase 1B. Nat. Commun. 11:788.

Humphrey, W., A. Dalke, and K. Schulten. 1996. VMD: visual molecular dynamics. J. Mol. Graph. 14:33–8, 27–8.

Izadi, S., R. Anandakrishnan, and A.V. Onufriev. 2014. Building water models, A different approach. arXiv [physics.chem-ph].

Jaroszewicz, W., J. Morcinek-Orłowska, K. Pierzynowska, L. Gaffke, and G. Węgrzyn. 2022. Phage display and other peptide display technologies. FEMS Microbiol. Rev. 46:fuab052.

Kawano, F., Y. Aono, H. Suzuki, and M. Sato. 2013. Fluorescence imaging-based high-throughput screening of fast- and slow-cycling LOV proteins. PLoS One. 8:e82693.

Klein, A.M., R.M. de Queiroz, D. Venkatesh, and C. Prives. 2021. The roles and regulation of MDM2 and MDMX: it is not just about p53. Genes Dev. 35:575–601.

Kubbutat, M.H., S.N. Jones, and K.H. Vousden. 1997. Regulation of p53 stability by Mdm2. Nature. 387:299–303.

Li, C., M. Pazgier, C. Li, W. Yuan, M. Liu, G. Wei, W.-Y. Lu, and W. Lu. 2010. Systematic mutational analysis of peptide inhibition of the p53-MDM2/MDMX interactions. J. Mol. Biol. 398:200–213.

Li, X., N. Gohain, S. Chen, Y. Li, X. Zhao, B. Li, W.D. Tolbert, W. He, M. Pazgier, H. Hu, and W. Lu. 2021. Design of ultrahigh-affinity and dual-specificity peptide antagonists of MDM2 and MDMX for P53 activation and tumor suppression. Acta Pharm Sin B. 11:2655–2669.

Li, X.-L., R. Tei, M. Uematsu, and J.M. Baskin. 2024. Ultralow background membrane editors for spatiotemporal control of phosphatidic acid metabolism and signaling. ACS Cent. Sci. 10:543–554.

Lungu, O.I., R.A. Hallett, E.J. Choi, M.J. Aiken, K.M. Hahn, and B. Kuhlman. 2012. Designing photoswitchable peptides using the AsLOV2 domain. Chem. Biol. 19:507–517.

Manoilov, K.Y., V.V. Verkhusha, and D.M. Shcherbakova. 2021. A guide to the optogenetic regulation of endogenous molecules. Nat. Methods. 18:1027–1037.

Marchenko, N.D., A. Zaika, and U.M. Moll. 2000. Death Signal-induced Localization of p53 Protein to Mitochondria: A POTENTIAL ROLE IN APOPTOTIC SIGNALING*. J. Biol. Chem. 275:16202–16212.

Melero-Fernandez de Mera, R.M., L.-L. Li, A. Popinigis, K. Cisek, M. Tuittila, L. Yadav, A. Serva, and M.J. Courtney. 2017. A simple optogenetic MAPK inhibitor design reveals resonance between transcription-regulating circuitry and temporally-encoded inputs. Nat. Commun. 8:15017.

Mihara, M., S. Erster, A. Zaika, O. Petrenko, T. Chittenden, P. Pancoska, and U.M. Moll. 2003. p53 has a direct apoptogenic role at the mitochondria. Mol. Cell. 11:577–590.

Murakoshi, H., M.E. Shin, P. Parra-Bueno, E.M. Szatmari, A.C.E. Shibata, and R. Yasuda. 2017. Kinetics of endogenous CaMKII required for synaptic plasticity revealed by optogenetic kinase inhibitor. Neuron. 94:37–47.e5.

Newton, M.S., Y. Cabezas-Perusse, C.L. Tong, and B. Seelig. 2020. In vitro selection of peptides and proteins-advantages of mRNA display. ACS Synth. Biol. 9:181–190.

Niopek, D., D. Benzinger, J. Roensch, T. Draebing, P. Wehler, R. Eils, and B. Di Ventura. 2014. Engineering light-inducible nuclear localization signals for precise spatiotemporal control of protein dynamics in living cells. Nat. Commun. 5:4404.

Niopek, D., P. Wehler, J. Roensch, R. Eils, and B. Di Ventura. 2016. Optogenetic control of nuclear protein export. Nat. Commun. 7:10624.

Nitiss, J.L. 2009. Targeting DNA topoisomerase II in cancer chemotherapy. Nat. Rev. Cancer. 9:338–350.

Oliner, J.D., J.A. Pietenpol, S. Thiagalingam, J. Gyuris, K.W. Kinzler, and B. Vogelstein. 1993. Oncoprotein MDM2 conceals the activation domain of tumour suppressor p53. Nature. 362:857–860.

Paonessa, F., S. Criscuolo, S. Sacchetti, D. Amoroso, H. Scarongella, F. Pecoraro Bisogni, E. Carminati, G. Pruzzo, L. Maragliano, F. Cesca, and F. Benfenati. 2016. Regulation of neural gene transcription by optogenetic inhibition of the RE1-silencing transcription factor. Proc. Natl. Acad. Sci. U. S. A. 113:E91–100.

Pazgier, M., M. Liu, G. Zou, W. Yuan, C. Li, C. Li, J. Li, J. Monbo, D. Zella, S.G. Tarasov, and W. Lu. 2009. Structural basis for high-affinity peptide inhibition of p53 interactions with MDM2 and MDMX. Proc. Natl. Acad. Sci. U. S. A. 106:4665–4670.

Phillips, A., A. Teunisse, S. Lam, K. Lodder, M. Darley, M. Emaduddin, A. Wolf, J. Richter, J. de Lange, M. Verlaan-de Vries, K. Lenos, A. Böhnke, F. Bartel, J.P. Blaydes, and A.G. Jochemsen. 2010. HDMX-L is expressed from a functional p53-responsive promoter in the first intron of the HDMX gene and participates in an autoregulatory feedback loop to control p53 activity. J. Biol. Chem. 285:29111–29127.

Qing, G., L.-C. Ma, A. Khorchid, G.V.T. Swapna, T.K. Mal, M.M. Takayama, B. Xia, S. Phadtare, H. Ke, T. Acton, G.T. Montelione, M. Ikura, and M. Inouye. 2004. Cold-shock induced high-yield protein production in Escherichia coli. Nat. Biotechnol. 22:877–882.

Quiroz, C., Y.B. Saavedra, B. Armijo-Galdames, J. Amado-Hinojosa, Á. Olivera-Nappa, A. Sanchez-Daza, and D. Medina-Ortiz. 2021. Peptipedia: a user-friendly web application and a comprehensive database for peptide research supported by Machine Learning approach. Database (Oxford). 2021:baab055.

Redchuk, T.A., E.S. Omelina, K.G. Chernov, and V.V. Verkhusha. 2017. Near-infrared optogenetic pair for protein regulation and spectral multiplexing. Nat. Chem. Biol. 13:633–639.

Repina, N.A., A. Rosenbloom, A. Mukherjee, D.V. Schaffer, and R.S. Kane. 2017. At light speed: Advances in optogenetic systems for regulating cell signaling and behavior. Annu. Rev. Chem. Biomol. Eng. 8:13–39.

Ryckaert, J.-P., G. Ciccotti, and H.J.C. Berendsen. 1977. Numerical integration of the cartesian equations of motion of a system with constraints: molecular dynamics of n-alkanes. J. Comput. Phys. 23:327–341.

Salomon, M., J.M. Christie, E. Knieb, U. Lempert, and W.R. Briggs. 2000. Photochemical and mutational analysis of the FMN-binding domains of the plant blue light receptor, phototropin. Biochemistry. 39:9401–9410.

Schindelin, J., I. Arganda-Carreras, E. Frise, V. Kaynig, M. Longair, T. Pietzsch, S. Preibisch, C. Rueden, S. Saalfeld, B. Schmid, J.-Y. Tinevez, D.J. White, V. Hartenstein, K. Eliceiri, P. Tomancak, and A. Cardona. 2012. Fiji: an open-source platform for biological-image analysis. Nat. Methods. 9:676–682.

Schmidt, U., M. Weigert, C. Broaddus, and G. Myers. 2018. Cell detection with star-convex polygons. In Medical Image Computing and Computer Assisted Intervention – MICCAI 2018. Springer International Publishing, Cham. 265–273.

Schon, O., A. Friedler, M. Bycroft, S.M.V. Freund, and A.R. Fersht. 2002. Molecular mechanism of the interaction between MDM2 and p53. J. Mol. Biol. 323:491–501.

Sengupta, A., Z. Li, L.F. Song, P. Li, and K.M. Merz Jr. 2021. Parameterization of monovalent ions for the OPC3, OPC, TIP3P-FB, and TIP4P-FB water models. J. Chem. Inf. Model. 61:869–880.

Shvarts, A., W.T. Steegenga, N. Riteco, T. van Laar, P. Dekker, M. Bazuine, R.C. van Ham, W. van der Houven van Oordt, G. Hateboer, A.J. van der Eb, and A.G. Jochemsen. 1996. MDMX: a novel p53-binding protein with some functional properties of MDM2. EMBO J. 15:5349–5357.

Strickland, D., Y. Lin, E. Wagner, C.M. Hope, J. Zayner, C. Antoniou, T.R. Sosnick, E.L. Weiss, and M. Glotzer. 2012. TULIPs: tunable, light-controlled interacting protein tags for cell biology. Nat. Methods. 9:379–384.

Tian, C., K. Kasavajhala, K.A.A. Belfon, L. Raguette, H. Huang, A.N. Migues, J. Bickel, Y. Wang, J. Pincay, Q. Wu, and C. Simmerling. 2020. Ff19SB: Amino-acid-specific protein backbone parameters trained against quantum mechanics energy surfaces in solution. J. Chem. Theory Comput. 16:528–552.

Tsuruoka, T., Y. Goto, and K. Aoki. 2025. Opto-p53: A light-controllable activation of p53 signaling pathway. Cell Struct. Funct. 25017.

Tsuruoka, T., E. Nakayama, T. Endo, S. Harashima, R. Kamada, K. Sakaguchi, and T. Imagawa. 2023. Development of a fluorescence reporter system to quantify transcriptional activity of endogenous p53 in living cells. J. Cell Sci. 136. doi:10.1242/jcs.260918.

Uda, Y., Y. Goto, S. Oda, T. Kohchi, M. Matsuda, and K. Aoki. 2017. Efficient synthesis of phycocyanobilin in mammalian cells for optogenetic control of cell signaling. Proc. Natl. Acad. Sci. U. S. A. 114:11962–11967.

Vassilev, L.T., B.T. Vu, B. Graves, D. Carvajal, F. Podlaski, Z. Filipovic, N. Kong, U. Kammlott, C. Lukacs, C. Klein, N. Fotouhi, and E.A. Liu. 2004. In vivo activation of the p53 pathway by small-molecule antagonists of MDM2. Science. 303:844–848.

Wade, M., Y.-C. Li, and G.M. Wahl. 2013. MDM2, MDMX and p53 in oncogenesis and cancer therapy. Nat. Rev. Cancer. 13:83–96.

Wagner, A.J., J.M. Kokontis, and N. Hay. 1994. Myc-mediated apoptosis requires wild-type p53 in a manner independent of cell cycle arrest and the ability of p53 to induce p21waf1/cip1. Genes Dev. 8:2817–2830.

Wang, H., M. Vilela, A. Winkler, M. Tarnawski, I. Schlichting, H. Yumerefendi, B. Kuhlman, R. Liu, G. Danuser, and K.M. Hahn. 2016. LOVTRAP: an optogenetic system for photoinduced protein dissociation. Nat. Methods. 13:755–758.

Wehler, P., and B. Di Ventura. 2019. Engineering Optogenetic Control of Endogenous p53 Protein Levels. NATO Adv. Sci. Inst. Ser. E Appl. Sci. 9:2095.

Wu, Y.I., D. Frey, O.I. Lungu, A. Jaehrig, I. Schlichting, B. Kuhlman, and K.M. Hahn. 2009. A genetically encoded photoactivatable Rac controls the motility of living cells. Nature. 461:104–108.

Yamamoto, K., H. Miura, M. Ishida, Y. Mii, N. Kinoshita, S. Takada, N. Ueno, S. Sawai, Y. Kondo, and K. Aoki. 2021. Optogenetic relaxation of actomyosin contractility uncovers mechanistic roles of cortical tension during cytokinesis. Nat. Commun. 12:7145.

Yang, R., B. Huang, Y. Zhu, Y. Li, F. Liu, and J. Shi. 2018. Cell type-dependent bimodal p53 activation engenders a dynamic mechanism of chemoresistance. Sci Adv. 4:eaat5077.

Yao, X., M.K. Rosen, and K.H. Gardner. 2008. Estimation of the available free energy in a LOV2-J alpha photoswitch. Nat. Chem. Biol. 4:491–497.

Yi, J.J., H. Wang, M. Vilela, G. Danuser, and K.M. Hahn. 2014. Manipulation of endogenous kinase activity in living cells using photoswitchable inhibitory peptides. ACS Synth. Biol. 3:788–795.

Yizhar, O., L.E. Fenno, T.J. Davidson, M. Mogri, and K. Deisseroth. 2011. Optogenetics in neural systems. Neuron. 71:9–34.

Yumerefendi, H., D.J. Dickinson, H. Wang, S.P. Zimmerman, J.E. Bear, B. Goldstein, K. Hahn, and B. Kuhlman. 2015. Control of Protein Activity and Cell Fate Specification via Light-Mediated Nuclear Translocation. PLoS One. 10:e0128443.

Yumerefendi, H., A.M. Lerner, S.P. Zimmerman, K. Hahn, J.E. Bear, B.D. Strahl, and B. Kuhlman. 2016. Light-induced nuclear export reveals rapid dynamics of epigenetic modifications. Nat. Chem. Biol. 12:399–401.

Yumerefendi, H., H. Wang, D.J. Dickinson, A.M. Lerner, P. Malkus, B. Goldstein, K. Hahn, and B. Kuhlman. 2018. Light-dependent cytoplasmic recruitment enhances the dynamic range of a nuclear import photoswitch. Chembiochem. 19:1319–1325.

Yusa, K., R. Rad, J. Takeda, and A. Bradley. 2009. Generation of transgene-free induced pluripotent mouse stem cells by the piggyBac transposon. Nat. Methods. 6:363–369.

Yusa, K., L. Zhou, M.A. Li, A. Bradley, and N.L. Craig. 2011. A hyperactive piggyBac transposase for mammalian applications. Proc. Natl. Acad. Sci. U. S. A. 108:1531–1536.

Zauberman, A., D. Flusberg, Y. Haupt, Y. Barak, and M. Oren. 1995. A functional p53-responsive intronic promoter is contained within the human mdm2 gene. Nucleic Acids Res. 23:2584–2592.

Zayner, J.P., C. Antoniou, and T.R. Sosnick. 2012. The amino-terminal helix modulates light-activated conformational changes in AsLOV2. J. Mol. Biol. 419:61–74.

Zhu, L., H.M. McNamara, and J.E. Toettcher. 2023. Light-switchable transcription factors obtained by direct screening in mammalian cells. Nat. Commun. 14:3185.

