## Supplementary Information for "Opto-MDMi: a dual-lock optogenetic system for robust activation of endogenous p53"

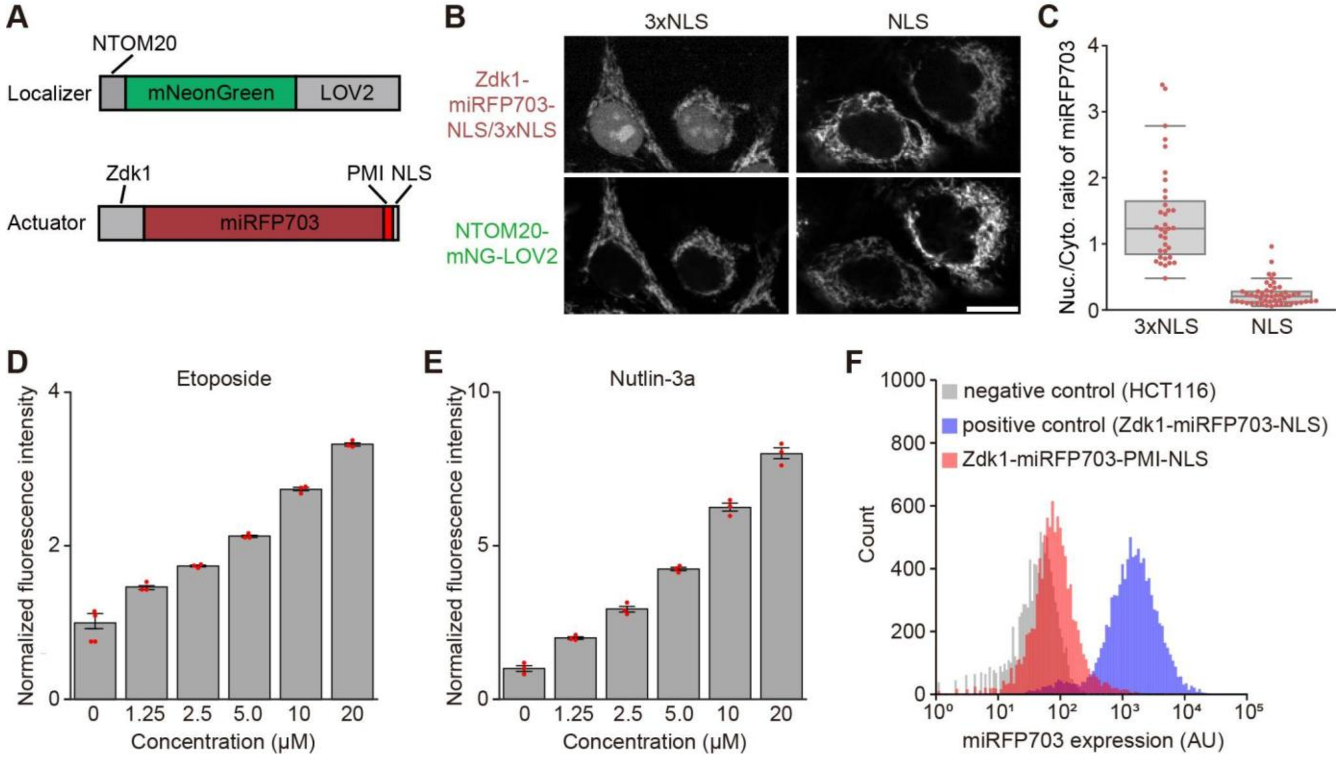


**Figure S1. Characterization of the Opto-MDMi (LOVTRAP) and p53 reporter system.**

A. Details of the expression constructs for the Opto-MDMi (LOVTRAP) localizer and actuator.
B. Subcellular localization of the actuator fragment with different NLS strengths (Zdk1-miRFP703-NLS or 3xNLS) and the localizer.
C. The ratio of the nuclear to cytosolic actuator signals under the condition of Fig. S1B.
D. Dose-response of the stable cell line harboring the p53 transcriptional reporter, measured 24 hours after treatment with the indicated concentrations of etoposide.
E. Dose-response of the stable cell line harboring the p53 transcriptional reporter, measured 24 hours after treatment with the indicated concentrations of nutlin-3a.
F. Distribution of the Opto-MDMi (LOVTRAP) actuator expression quantified by flow cytometry. Bulk stable cell lines expressing the p53 transcriptional reporter, the Opto-MDMi (LOVTRAP) localizer, and the actuator were used. Expression of the Opto-MDMi (LOVTRAP) actuator was quantified by the miRFP703 fluorescence signal.

**A**

```
Name Sequence
PMI truncation
WT    LOV2(404-546)-sr-TSFAEYWNLLSP
-1N   LOV2(404-546)-sr- SFAEYWNLLSP
-2N   LOV2(404-546)-sr-  FAEYWNLLSP
-3N   LOV2(404-546)-sr-   AEYWNLLSP
-1C   LOV2(404-546)-sr-TSFAEYWNLLS
-2C   LOV2(404-546)-sr-TSFAEYWNLL
-3C   LOV2(404-546)-sr-TSFAEYWNL

PMI-M3 truncation
WT    LOV2(404-546)-sr-LTFLEYWAQLMQ
-1N   LOV2(404-546)-sr- TFLEYWAQLMQ
-2N   LOV2(404-546)-sr-  FLEYWAQLMQ
-3N   LOV2(404-546)-sr-   LEYWAQLMQ
-1C   LOV2(404-546)-sr-LTFLEYWAQLM
-2C   LOV2(404-546)-sr-LTFLEYWAQL
-3C   LOV2(404-546)-sr-LTFLEYWAQ
```

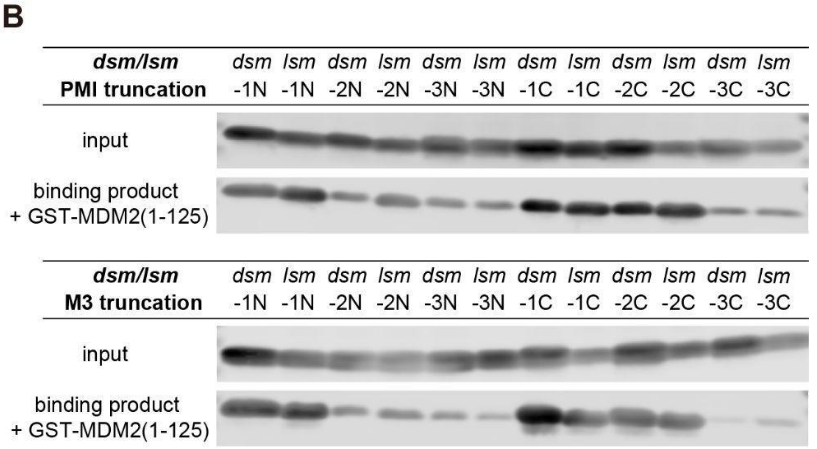


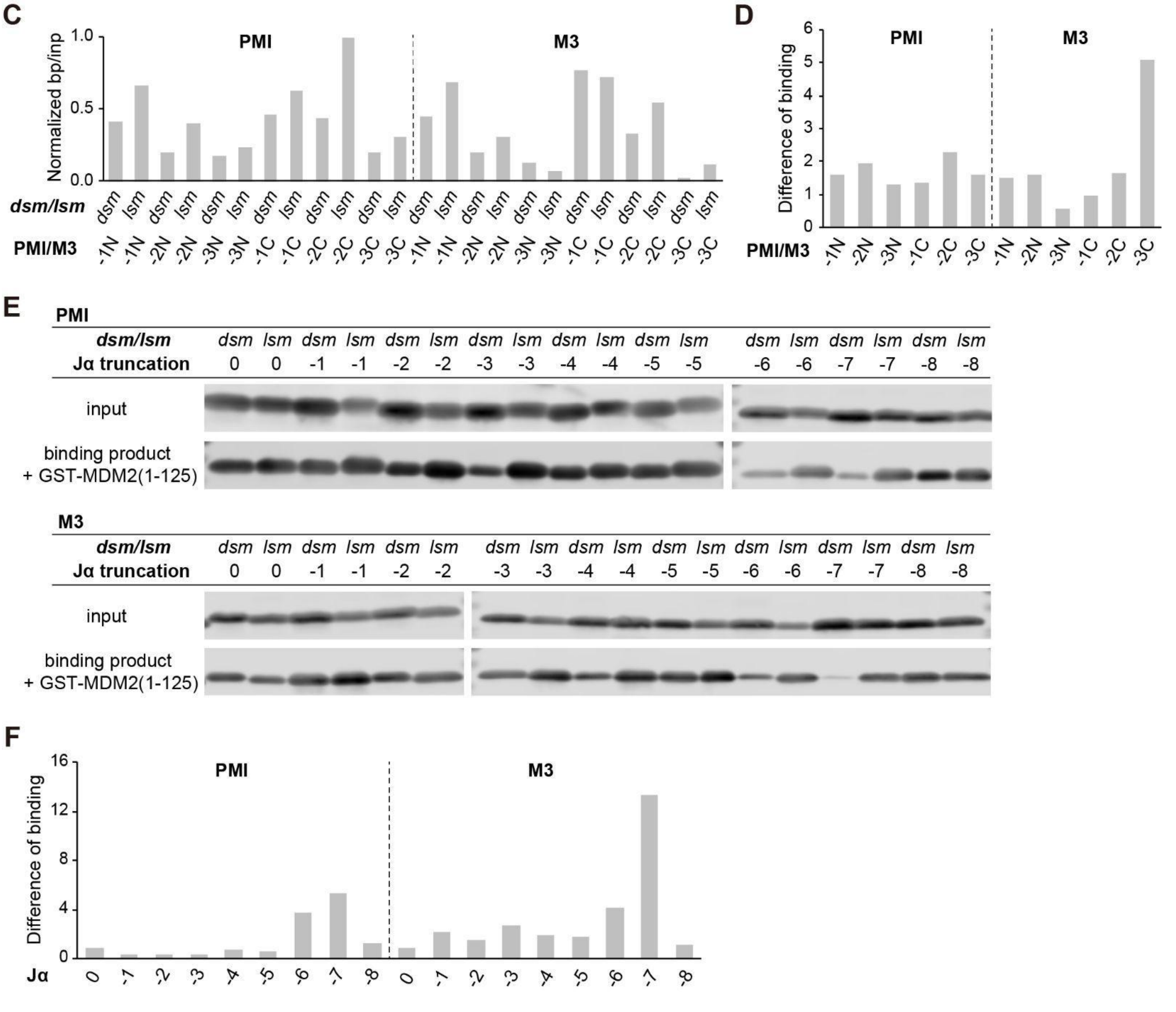

**Figure S2. Results of *in vitro* screening using expression libraries with PMI or Jα helix truncations.**

A. The expression library of the LOV2-PMI/PMI-M3 fragments with different PMI/PMI-M3 truncations used in the *in vitro* binding assay.
B. Western blot images detecting each LOV2-PMI/PMI-M3 fragment with different PMI/PMI-M3 truncations bound to GST-MDM2(1–125).
C. Ratio of the signal intensities between the input and binding products shown in Fig. S2B.
D. Differences in binding activity between the *dsm* or *lsm* of each LOV2-PMI/PMI-M3 fragment shown in Fig. S2B.
E. Western blot images detecting each LOV2-PMI/PMI-M3 fragment with different Jα-helix truncation bound to GST-MDM2(1–125).
F. Differences in binding activity between the *dsm* or *lsm* of each LOV2-PMI/PMI-M3 fragments shown in Fig. S2E.

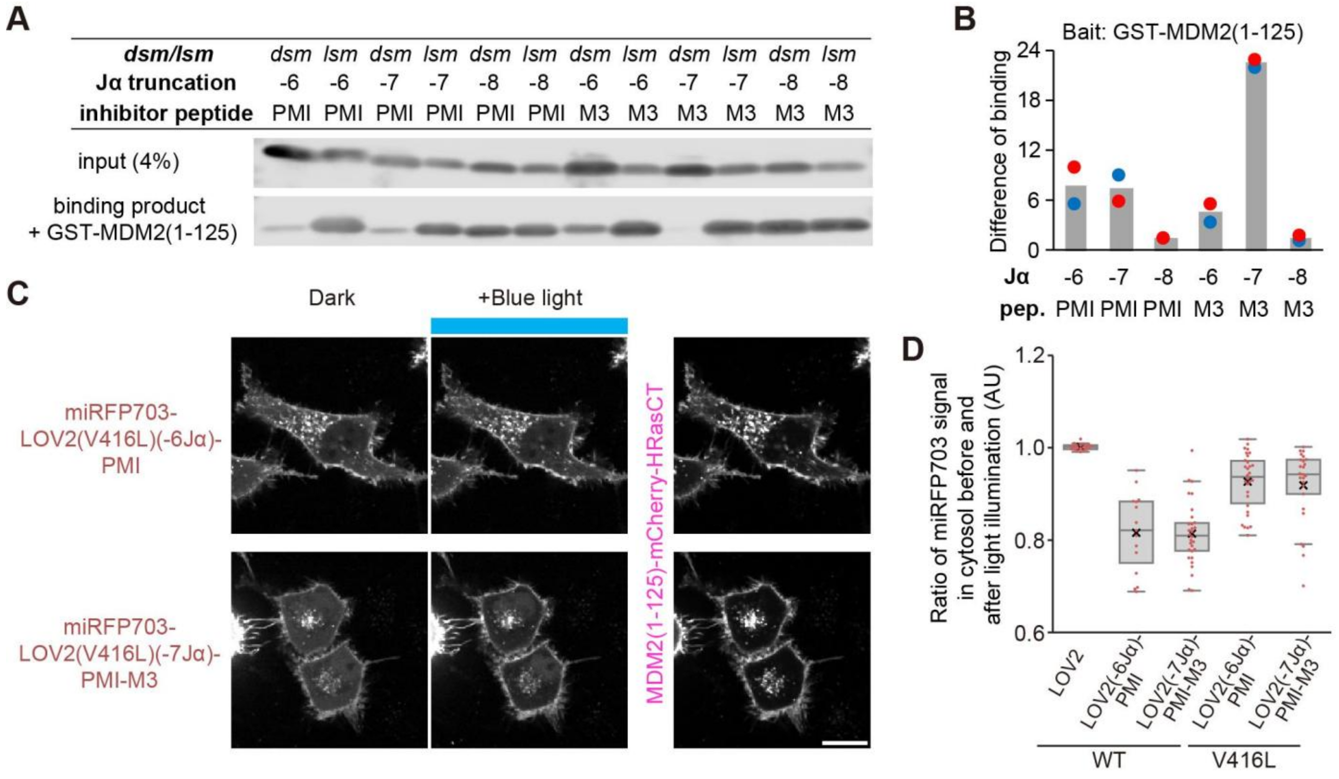


**Figure S3. Evaluation of the LOV2(V416L)-PMI modules *in vitro* and in cultured cells.**

A. Western blot images detecting each LOV2(V416L)-PMI/PMI-M3 fragment with different PMI/PMI-M3 truncations bound to GST-MDM2(1–125).

B. Differences in binding activity between the *dsm* or *lsm* of each LOV2(V416L)-PMI/PMI-M3 fragment shown in Fig. S3A. Blue and red points indicate experimental replicates.

C. Light-dependent change in the localization of the actuator fragment harboring the LOV2(V416L) mutation. Scale bar: 20 μm.

D. Changes in the cytosolic actuator levels before and after the blue-light illumination. Values were calculated by dividing the cytosolic actuator signal after light illumination by the signal before light illumination.

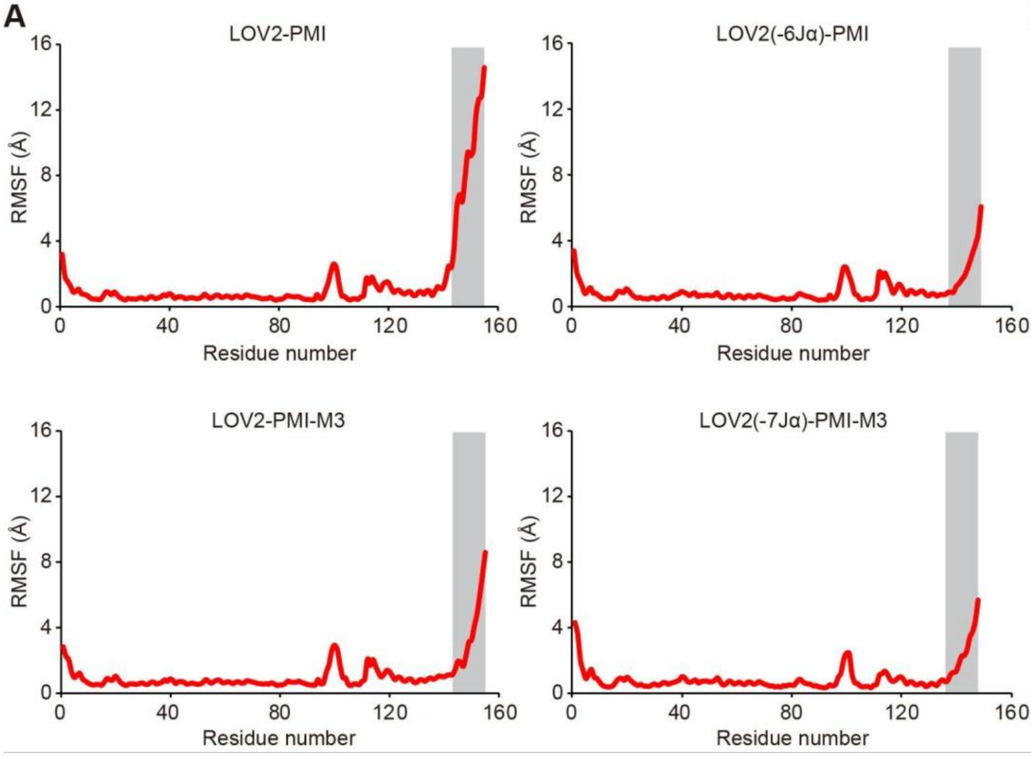


**B** SASA (Å²)

| PMI # | AA | LOV2-PMI | LOV2(-6Jα)-PMI |
|---|---|---|---|
| 1 | Thr | 70 | 75 |
| 2 | Ser | 65 | 12 |
| 3 | Phe | 58 | 10 |
| 4 | Ala | 45 | 54 |
| 5 | Glu | 126 | 96 |
| 6 | Tyr | 125 | 43 |
| 7 | Trp | 97 | 97 |
| 8 | Asn | 105 | 102 |
| 9 | Leu | 151 | 141 |
| 10 | Leu | 97 | 102 |
| 11 | Ser | 66 | 60 |
| 12 | Pro | 177 | 191 |

| PMI-M3 # | AA | LOV2-PMI-M3 | LOV2(-7Jα)-PMI-M3 |
|---|---|---|---|
| 1 | Leu | 26 | 38 |
| 2 | Thr | 68 | 76 |
| 3 | Phe | 76 | 54 |
| 4 | Leu | 65 | 1 |
| 5 | Glu | 74 | 102 |
| 6 | Tyr | 112 | 129 |
| 7 | Trp | 135 | 57 |
| 8 | Ala | 69 | 26 |
| 9 | Gln | 117 | 118 |
| 10 | Leu | 116 | 115 |
| 11 | Met | 154 | 100 |
| 12 | Gln | 189 | 191 |

**Figure S4. Calculated RMSF and SASA values from the molecular dynamics simulations.**

A. Calculated root-mean-square fluctuation (RMSF) values of the Cα atoms in all residues. The grey area indicates the residues of the PMI/PMI-M3 peptides.

B. Calculated solvent-accessible surface area (SASA) values for all residues in the PMI/PMI-M3 peptides. The color intensity of the table was varied according to the SASA values.

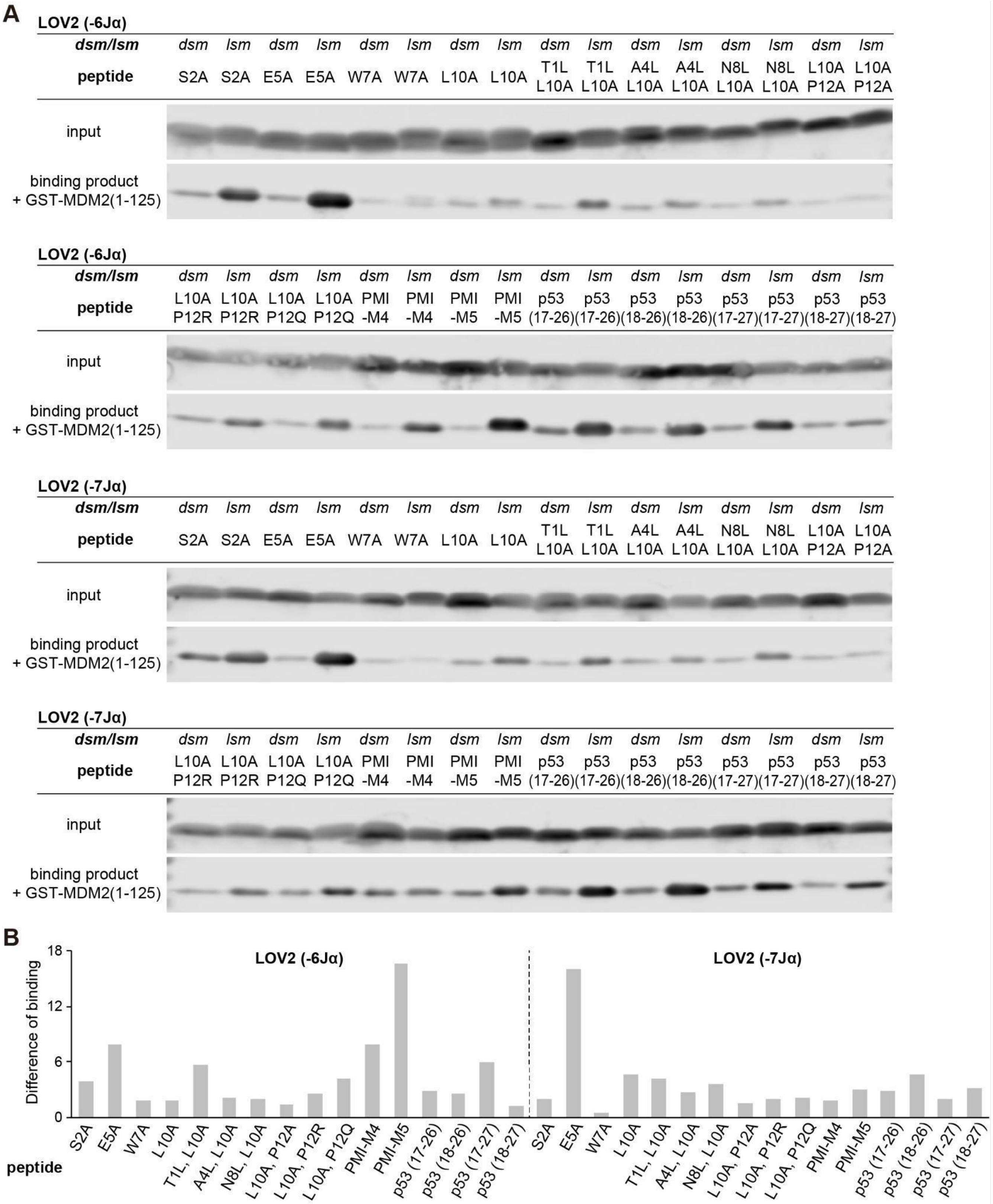
A
LOV2 (-6Jα)
dsm/lsm
dsm lsm dsm lsm dsm lsm dsm lsm dsm lsm dsm lsm dsm lsm dsm lsm
peptide
S2A S2A E5A E5A W7A W7A L10A L10A T1L L10A T1L L10A A4L L10A A4L L10A N8L L10A N8L L10A L10A P12A L10A P12A
input
binding product + GST-MDM2(1-125)
LOV2 (-6Jα)
dsm/lsm
dsm lsm dsm lsm dsm lsm dsm lsm dsm lsm dsm lsm dsm lsm dsm lsm
peptide
L10A P12R L10A P12R L10A P12Q L10A P12Q PMI -M4 PMI -M4 PMI -M5 PMI -M5 p53 (17-26) p53 (17-26) p53 (18-26) p53 (18-26) p53 (17-27) p53 (17-27) p53 (18-27) p53 (18-27)
input
binding product + GST-MDM2(1-125)
LOV2 (-7Jα)
dsm/lsm
dsm lsm dsm lsm dsm lsm dsm lsm dsm lsm dsm lsm dsm lsm dsm lsm
peptide
S2A S2A E5A E5A W7A W7A L10A L10A T1L L10A T1L L10A A4L L10A A4L L10A N8L L10A N8L L10A L10A P12A L10A P12A
input
binding product + GST-MDM2(1-125)
LOV2 (-7Jα)
dsm/lsm
dsm lsm dsm lsm dsm lsm dsm lsm dsm lsm dsm lsm dsm lsm dsm lsm
peptide
L10A P12R L10A P12R L10A P12Q L10A P12Q PMI -M4 PMI -M4 PMI -M5 PMI -M5 p53 (17-26) p53 (17-26) p53 (18-26) p53 (18-26) p53 (17-27) p53 (17-27) p53 (18-27) p53 (18-27)
input
binding product + GST-MDM2(1-125)
B
Difference of binding
0 6 12 18
LOV2 (-6Jα)
LOV2 (-7Jα)
peptide
S2A E5A W7A L10A T1L, L10A A4L, L10A N8L, L10A L10A, P12A L10A, P12R L10A, P12Q PMI-M4 PMI-M5 p53 (17-26) p53 (18-26) p53 (17-27) p53 (18-27)
S2A E5A W7A L10A T1L, L10A A4L, L10A N8L, L10A L10A, P12A L10A, P12R L10A, P12Q PMI-M4 PMI-M5 p53 (17-26) p53 (18-26) p53 (17-27) p53 (18-27)

**Figure S5. Results of *in vitro* screening using an expression library with various PMI peptides.**

A. Western blot images detecting each LOV2-inhibitory peptide fragment with different sequences bound to GST-MDM2(1–125).
B. Differences in binding activity between the *dsm* or *lsm* of each LOV2-PMI/PMI-M3 fragment shown in Fig. S5A.

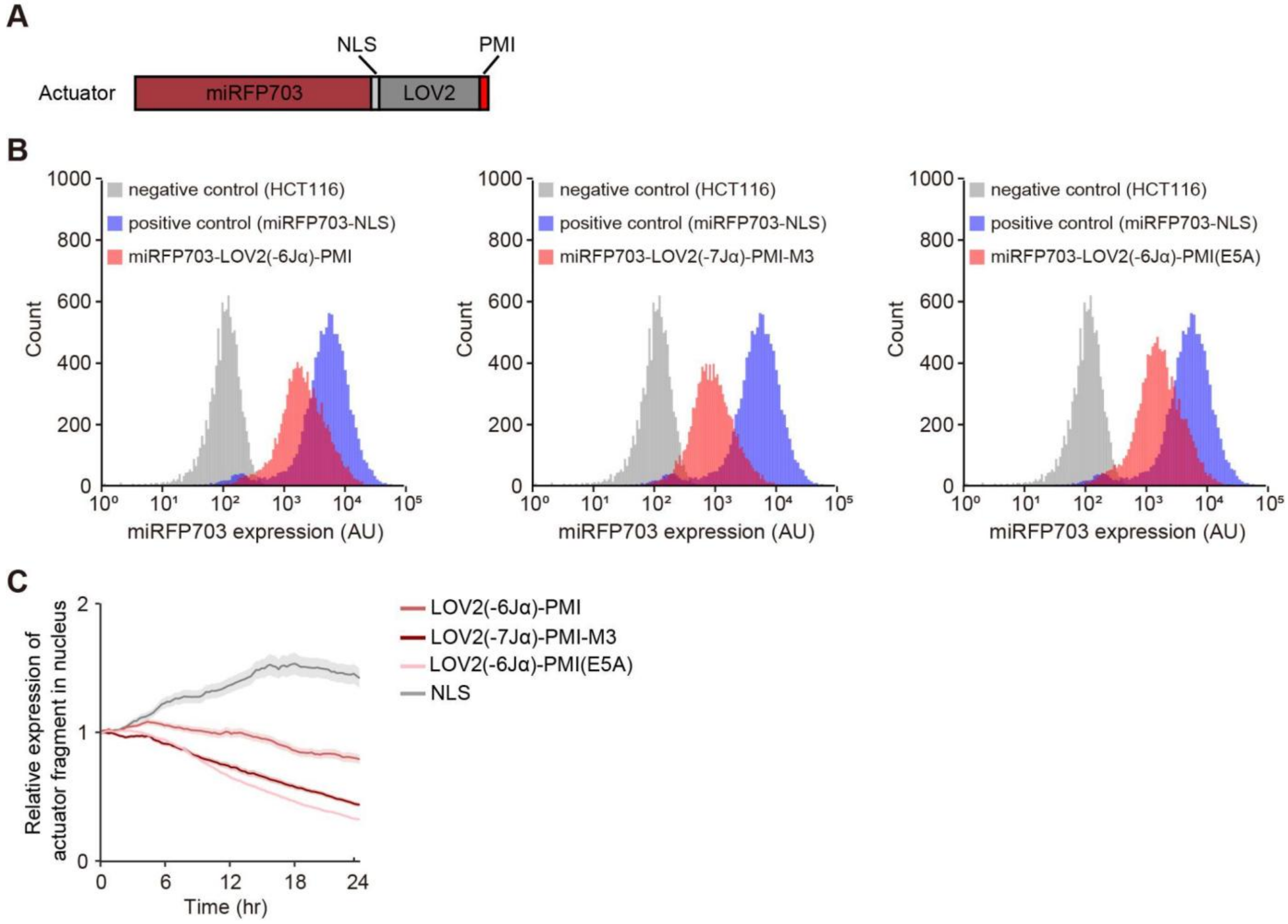


**Figure S6. Characterization of the Opto-MDMi (LOV2-PMI) system.**

A. Details of the expression constructs for the Opto-MDMi (LOV2-PMI) actuator.
B. Distribution of Opto-MDMi (LOV2-PMI) actuator expression quantified by flow cytometry. Bulk stable cell lines expressing the p53 transcriptional reporter, the Opto-MDMi (LOV2-PMI) localizer, and the actuator were used. Expression of the Opto-MDMi (LOV2-PMI) actuator was quantified by the miRFP703 fluorescence signal.
C. Temporal changes in the nuclear expression of the Opto-MDMi (LOV2-PMI) actuator in each cell line. The plot shows the mean ± s.e.m. The cell trace data are the same as those shown in Fig. 6D.

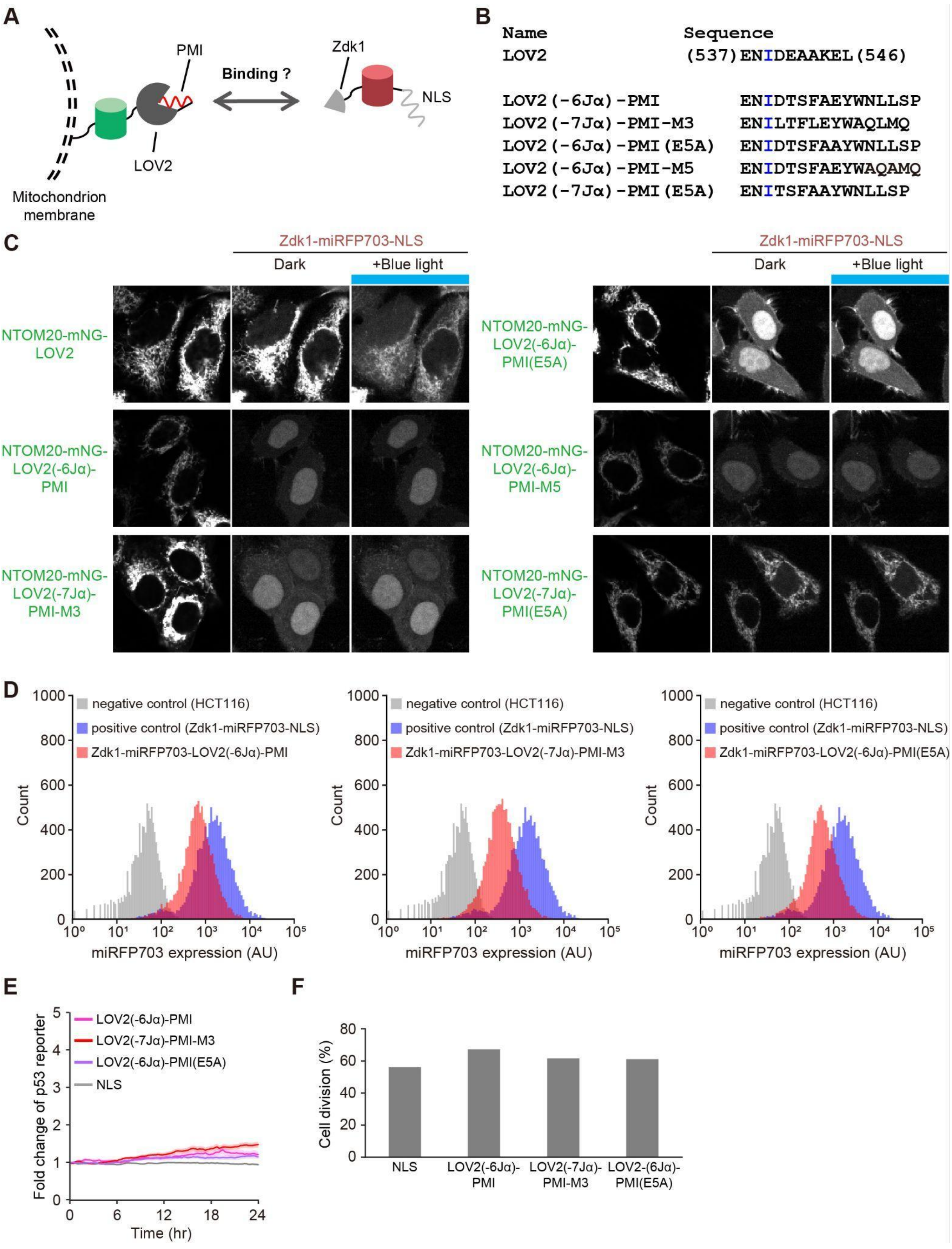
A
PMI
Binding ?
Zdk1
NLS
LOV2
Mitochondrion membrane
B
Name Sequence
LOV2 (537)ENIDEAAKEL(546)
LOV2(-6Jα)-PMI ENIDTSFAEYWNLLSP
LOV2(-7Jα)-PMI-M3 ENILTFLEYWAQLMQ
LOV2(-6Jα)-PMI(E5A) ENIDTSFAAYWNLLSP
LOV2(-6Jα)-PMI-M5 ENIDTSFAEYWAQAMQ
LOV2(-7Jα)-PMI(E5A) ENITSFAAYWNLLSP
C
Zdk1-miRFP703-NLS
Dark
+Blue light
NTOM20-mNG-LOV2
NTOM20-mNG-LOV2(-6Jα)-PMI
NTOM20-mNG-LOV2(-7Jα)-PMI-M3
NTOM20-mNG-LOV2(-6Jα)-PMI(E5A)
NTOM20-mNG-LOV2(-6Jα)-PMI-M5
NTOM20-mNG-LOV2(-7Jα)-PMI(E5A)
D
negative control (HCT116)
positive control (Zdk1-miRFP703-NLS)
Zdk1-miRFP703-LOV2(-6Jα)-PMI
Zdk1-miRFP703-LOV2(-7Jα)-PMI-M3
Zdk1-miRFP703-LOV2(-6Jα)-PMI(E5A)
Count
miRFP703 expression (AU)
E
LOV2(-6Jα)-PMI
LOV2(-7Jα)-PMI-M3
LOV2(-6Jα)-PMI(E5A)
NLS
Fold change of p53 reporter
Time (hr)
F
Cell division (%)
NLS
LOV2(-6Jα)-PMI
LOV2(-7Jα)-PMI-M3
LOV2-(6Jα)-PMI(E5A)

**Figure S7. Characterization of the Opto-MDMi system.**

A. A schematic illustration of the interaction assay between the Opto-MDMi (LOV2-PMI) actuator and the Zdk1 fragment.
B. The expression constructs developed for the interaction assay.
C. Subcellular localization of the Zdk1-miRFP703-NLS fragment when co-expressed with each mitochondria-tethered Opto-MDMi (LOV2-PMI) actuator fragment.
D. Distribution of Opto-MDMi actuator expression quantified by flow cytometry. Bulk stable cell lines expressing the p53 transcriptional reporter, the Opto-MDMi localizer, and the actuator were used. Expression of the Opto-MDMi actuator was quantified by the miRFP703 fluorescence signal.
E. Temporal changes in the p53 transcription reporter in each cell line. The plot shows the mean ± s.e.m. (LOV2(–6Jα)-PMI, n = 128 cells; LOV2(–7Jα)-PMI-M3, n = 117 cells; LOV2(–6Jα)-PMI(E5A), n = 118 cells; NLS, n = 164 cells).
F. Quantification of cell division events during the period from 0 to 24 hours in panel E.

### Table S1

| Plasmid name | Figures | Source or reference | Sequence |
|---|---|---|---|
| pCSIIneo-H1-mCherry | 1 | (Tany et al. 2022) | https://benchling.com/s/seq-IXstAoN97g80WlfDhgns |
| pPBbsr2-NTOM20-lin-mNeonGreen-lin-LOV2 | 1, 7, S1, S6 | This study | https://benchling.com/s/seq-5s2FjKIEXoTTQyCp1D08?m=slm-HwOhSe3BLz6V5oeob4ys |
| pPBneo-Zdk1-lin-miRFP703-dSal-NLS | 1, 6, 7, S1, S6, S7 | This study | https://benchling.com/s/seq-qwFJRnYRm7yL1XcujIzN?m=slm-LiuREz7A7UBwQf3jx406 |
| pPBneo-Zdk1-lin-miRFP703-dSal-NLSx3 | S1 | This study | https://benchling.com/s/seq-aKrLjkxG6ORZGnzG1Iyn?m=slm-ZMxI5Jor415kwcagrEzC |
| pPB-p53RE(CDKN1A)-CMVmin-mScarlet-I-NLSx3-AU1-PGKpuro | 1, 6, 7, S1 | (Tsuruoka et al. 2025) | https://benchling.com/s/seq-lv1G3m5olUoQKYBtBMca?m=slm-CbOooCTkrhJDY4fPlsMi |
| pPBneo-Zdk1-lin-miRFP703-dSal-PMI-NLS | 1, S1 | This study | https://benchling.com/s/seq-qamPGeoWYETiq7xfogcR?m=slm-vUjqqglCNkA4EWInbmH6 |
| pCold-GST-MDM2(1-125) | 2, 5, S2, S3, S5 | This study | https://benchling.com/s/seq-4ebIWyWIyTrEmXcYFj5y?m=slm-5PXD17yGWQW1rTvVmzNc |
| pCold-GST-MDMX(1-125) | 2, 5 | This study | https://benchling.com/s/seq-truDZAsnB8pFhMTo9sYP?m=slm-UyWPrsQydfQjCnqz59Bu |
| pCold-GST | 2 | This study | https://benchling.com/s/seq-OvhVYBPr6KYjAsZ9ozRH |
| pCAGGS-MDM2(1-125)-lin-mCherry-HRasCT | 3, 5, S3 | This study | https://benchling.com/s/seq-Pibo7oSmVmTHnKMACoSA?m=slm-mjxe4hpFNCmfzpsWTWYd |
| pCAGGS-miRFP703-dSal-lin-LOV2 | 3, S3 | This study | https://benchling.com/s/seq-3KFStEbOO0EW1L0uI7kO?m=slm-KCQ0Y2ULhPr2SbeaokSY |
| pCAGGS-miRFP703-dSal-lin-LOV2(-6Jα)-PMI | 3, S3 | This study | https://benchling.com/s/seq-asLu7wHSOxwzz54r5Cin?m=slm-zGKq10Kx86ZtbyCrV58D |
| pCAGGS-miRFP703-dSal-lin-LOV2(-7Jα)-PMI-M3 | 3, S3 | This study | https://benchling.com/s/seq-BUQT04r6bHsKn4Oopyv3?m=slm-20Riz3EjD7AWHpZLUByd |
| pCAGGS-miRFP703-dSal-lin-LOV2(-8Jα)-PMI-M3 | 3 | This study | https://benchling.com/s/seq-QBcz9wu56VllCDJJaw6c?m=slm-Fbg4OUpWzW2LzJp67LkK |

| | | | |
|---|---|---|---|
| pCAGGS-miRFP703-dSal-lin-LOV2(V416L)(-6Jα)-PMI | S3 | This study | https://benchling.com/s/seq-d8EBzGVzszyhRgYVs3aX?m=slm-m8g65ebbqWEOh1eeWG7Z |
| pCAGGS-miRFP703-dSal-lin-LOV2(V416L)(-7Jα)-PMI-M3 | S3 | This study | https://benchling.com/s/seq-jM5ZaMVGKPJbUWHRtJEP?m=slm-vGvPke4QslT2NgibMZVe |
| pCAGGS-miRFP703-dSal-lin-LOV2(-6Jα)-PMI(E5A) | 5 | This study | https://benchling.com/s/seq-p171IzRGZIuut2l13ciO?m=slm-NOTOgMVeLoGhRGBLijjI |
| pCAGGS-miRFP703-dSal-lin-LOV2(-6Jα)-PMI-M5 | 5 | This study | https://benchling.com/s/seq-pQU5evq2N24CH2wvB749?m=slm-BBGt18Dx65rYepHuhwEM |
| pCAGGS-miRFP703-dSal-lin-LOV2(-7Jα)-PMI(E5A) | 5 | This study | https://benchling.com/s/seq-TusVmxbdxYwYcZvm4o1R?m=slm-QZapTOiLDpaLIznNn566 |
| pPBneo-Zdk1-lin-miRFP703-dSal-lin-NLS-lin-LOV2(-6Jα)-PMI | 6, 7, S6, S7 | This study | https://benchling.com/s/seq-Ucnt2IaczKtEBkEX2me7?m=slm-J0lTou1hhoP7mywLVvvi |
| pPBneo-Zdk1-lin-miRFP703-dSal-lin-NLS-lin-LOV2(-7Jα)-PMI-M3 | 6, 7, S6, S7 | This study | https://benchling.com/s/seq-42ib6bowVqdGWRDRAnaT?m=slm-I684jSau8En76E9vPtwr |
| pPBneo-Zdk1-lin-miRFP703-dSal-lin-NLS-lin-LOV2(-6Jα)-PMI(E5A) | 6, 7, S6, S7 | This study | https://benchling.com/s/seq-zN2f2E1Bbkh466KNVg3Z?m=slm-xJzE2PbrLE79Qc9hnGdq |
| pPBbsr2-NTOM20-lin-mNeonGreen-lin-LOV2(-6Jα)-PMI | S7 | This study | https://benchling.com/s/seq-HAOqYDBqXWmTYEjEfB9P?m=slm-4fxHSbwVG71L37FEtVAk |
| pPBbsr2-NTOM20-lin-mNeonGreen-lin-LOV2(-7Jα)-PMI-M3 | S7 | This study | https://benchling.com/s/seq-s0xFIvxHgrWXlcKrKY5d?m=slm-96BKPw9JtuL2VaxfzvvC |
| pPBbsr2-NTOM20-lin-mNeonGreen-lin-LOV2(-6Jα)-PMI(E5A) | S7 | This study | https://benchling.com/s/seq-PdkvbhIXgGbAfvtB8U01?m=slm-Ql94z5YfoRmkn5VSDsVm |
| pPBbsr2-NTOM20-lin-mNeonGreen-lin-LOV2(-6Jα)-PMI-M5 | S7 | This study | https://benchling.com/s/seq-68rOjcMDEwywQaVzCykP?m=slm-vmQbXVmgclARY7YAzgjd |
| pPBbsr2-NTOM20-lin-mNeonGreen-lin-LOV2(-7Jα)-PMI(E5A) | S7 | This study | https://benchling.com/s/seq-lNt3M6WHhJjgvykQHrRh?m=slm-yfWejfBF3NOwojq3Zcx5 |

**Table S1. List of plasmids used in this study.**

### Table S2

| Plasmid name | Figures | Source or reference |
|---|---|---|
| pCold-T7-lin-NLS-lin-LOV2(C450A)-PMI(-1N) | S2 | This study |
| pCold-T7-lin-NLS-lin-LOV2(C450A)-PMI(-2N) | S2 | This study |
| pCold-T7-lin-NLS-lin-LOV2(C450A)-PMI(-3N) | S2 | This study |
| pCold-T7-lin-NLS-lin-LOV2(C450A)-PMI(-1C) | S2 | This study |
| pCold-T7-lin-NLS-lin-LOV2(C450A)-PMI(-2C) | S2 | This study |
| pCold-T7-lin-NLS-lin-LOV2(C450A)-PMI(-3C) | S2 | This study |
| pCold-T7-lin-NLS-lin-LOV2(I539E)-PMI(-1N) | S2 | This study |
| pCold-T7-lin-NLS-lin-LOV2(I539E)-PMI(-2N) | S2 | This study |
| pCold-T7-lin-NLS-lin-LOV2(I539E)-PMI(-3N) | S2 | This study |
| pCold-T7-lin-NLS-lin-LOV2(I539E)-PMI(-1C) | S2 | This study |
| pCold-T7-lin-NLS-lin-LOV2(I539E)-PMI(-2C) | S2 | This study |
| pCold-T7-lin-NLS-lin-LOV2(I539E)-PMI(-3C) | S2 | This study |
| pCold-T7-lin-NLS-lin-LOV2(C450A)-PMI-M3(-1N) | S2 | This study |
| pCold-T7-lin-NLS-lin-LOV2(C450A)-PMI-M3(-2N) | S2 | This study |
| pCold-T7-lin-NLS-lin-LOV2(C450A)-PMI-M3(-3N) | S2 | This study |
| pCold-T7-lin-NLS-lin-LOV2(C450A)-PMI-M3(-1C) | S2 | This study |
| pCold-T7-lin-NLS-lin-LOV2(C450A)-PMI-M3(-2C) | S2 | This study |
| pCold-T7-lin-NLS-lin-LOV2(C450A)-PMI-M3(-3C) | S2 | This study |
| pCold-T7-lin-NLS-lin-LOV2(I539E)-PMI-M3(-1N) | S2 | This study |
| pCold-T7-lin-NLS-lin-LOV2(I539E)-PMI-M3(-2N) | S2 | This study |
| pCold-T7-lin-NLS-lin-LOV2(I539E)-PMI-M3(-3N) | S2 | This study |
| pCold-T7-lin-NLS-lin-LOV2(I539E)-PMI-M3(-1C) | S2 | This study |
| pCold-T7-lin-NLS-lin-LOV2(I539E)-PMI-M3(-2C) | S2 | This study |
| pCold-T7-lin-NLS-lin-LOV2(I539E)-PMI-M3(-3C) | S2 | This study |

**Table S2. List of plasmids list for the PMI-truncated LOV2-PMI expression library used in the *in vitro* binding assay.**

### Table S3

| Plasmid name | Figures | Source or reference |
|---|---|---|
| pCold-T7-lin-NLS-lin-LOV2(C450A)(0Jα)-PMI | S2 | This study |
| pCold-T7-lin-NLS-lin-LOV2(C450A)(-1Jα)-PMI | S2 | This study |
| pCold-T7-lin-NLS-lin-LOV2(C450A)(-2Jα)-PMI | S2 | This study |
| pCold-T7-lin-NLS-lin-LOV2(C450A)(-3Jα)-PMI | S2 | This study |
| pCold-T7-lin-NLS-lin-LOV2(C450A)(-4Jα)-PMI | S2 | This study |
| pCold-T7-lin-NLS-lin-LOV2(C450A)(-5Jα)-PMI | S2 | This study |
| pCold-T7-lin-NLS-lin-LOV2(C450A)(-6Jα)-PMI | 2, S2 | This study |
| pCold-T7-lin-NLS-lin-LOV2(C450A)(-7Jα)-PMI | 2, S2 | This study |
| pCold-T7-lin-NLS-lin-LOV2(C450A)(-8Jα)-PMI | 2, S2 | This study |
| pCold-T7-lin-NLS-lin-LOV2(I539E)(0Jα)-PMI | S2 | This study |
| pCold-T7-lin-NLS-lin-LOV2(I539E)(-1Jα)-PMI | S2 | This study |
| pCold-T7-lin-NLS-lin-LOV2(I539E)(-2Jα)-PMI | S2 | This study |
| pCold-T7-lin-NLS-lin-LOV2(I539E)(-3Jα)-PMI | S2 | This study |
| pCold-T7-lin-NLS-lin-LOV2(I539E)(-4Jα)-PMI | S2 | This study |
| pCold-T7-lin-NLS-lin-LOV2(I539E)(-5Jα)-PMI | S2 | This study |
| pCold-T7-lin-NLS-lin-LOV2(I539E)(-6Jα)-PMI | 2, S2 | This study |
| pCold-T7-lin-NLS-lin-LOV2(I539E)(-7Jα)-PMI | 2, S2 | This study |
| pCold-T7-lin-NLS-lin-LOV2(I539E)(-8Jα)-PMI | 2, S2 | This study |
| pCold-T7-lin-NLS-lin-LOV2(C450A)(0Jα)-PMI-M3 | S2 | This study |
| pCold-T7-lin-NLS-lin-LOV2(C450A)(-1Jα)-PMI-M3 | S2 | This study |
| pCold-T7-lin-NLS-lin-LOV2(C450A)(-2Jα)-PMI-M3 | S2 | This study |
| pCold-T7-lin-NLS-lin-LOV2(C450A)(-3Jα)-PMI-M3 | S2 | This study |
| pCold-T7-lin-NLS-lin-LOV2(C450A)(-4Jα)-PMI-M3 | S2 | This study |
| pCold-T7-lin-NLS-lin-LOV2(C450A)(-5Jα)-PMI-M3 | S2 | This study |
| pCold-T7-lin-NLS-lin-LOV2(C450A)(-6Jα)-PMI-M3 | 2, S2 | This study |
| pCold-T7-lin-NLS-lin-LOV2(C450A)(-7Jα)-PMI-M3 | 2, S2 | This study |
| pCold-T7-lin-NLS-lin-LOV2(C450A)(-8Jα)-PMI-M3 | 2, S2 | This study |
| pCold-T7-lin-NLS-lin-LOV2(I539E)(0Jα)-PMI-M3 | S2 | This study |
| pCold-T7-lin-NLS-lin-LOV2(I539E)(-1Jα)-PMI-M3 | S2 | This study |
| pCold-T7-lin-NLS-lin-LOV2(I539E)(-2Jα)-PMI-M3 | S2 | This study |
| pCold-T7-lin-NLS-lin-LOV2(I539E)(-3Jα)-PMI-M3 | S2 | This study |

| | | |
|---|---|---|
| pCold-T7-lin-NLS-lin-LOV2(I539E)(-4Jα)-PMI-M3 | S2 | This study |
| pCold-T7-lin-NLS-lin-LOV2(I539E)(-5Jα)-PMI-M3 | S2 | This study |
| pCold-T7-lin-NLS-lin-LOV2(I539E)(-6Jα)-PMI-M3 | 2, S2 | This study |
| pCold-T7-lin-NLS-lin-LOV2(I539E)(-7Jα)-PMI-M3 | 2, S2 | This study |
| pCold-T7-lin-NLS-lin-LOV2(I539E)(-8Jα)-PMI-M3 | 2, S2 | This study |
| pCold-T7-mEGFP-PMI | 2, 5 | This study |
| pCold-T7-mEGFP-PMI-M3 | 2 | This study |
| pCold-T7-mEGFP | 2, 5 | This study |

**Table S3. List of plasmids for the Jα helix-truncated LOV2-PMI expression library used in the *in vitro* binding assay.**

### Table S4

| Plasmid name | Figures | Source or reference |
|---|---|---|
| pCold-T7-lin-NLS-lin-LOV2(V416L, C450A)(-6Jα)-PMI | S3 | This study |
| pCold-T7-lin-NLS-lin-LOV2(V416L, C450A)(-7Jα)-PMI | S3 | This study |
| pCold-T7-lin-NLS-lin-LOV2(V416L, C450A)(-8Jα)-PMI | S3 | This study |
| pCold-T7-lin-NLS-lin-LOV2(V416L, I539E)(-6Jα)-PMI | S3 | This study |
| pCold-T7-lin-NLS-lin-LOV2(V416L, I539E)(-7Jα)-PMI | S3 | This study |
| pCold-T7-lin-NLS-lin-LOV2(V416L, I539E)(-8Jα)-PMI | S3 | This study |
| pCold-T7-lin-NLS-lin-LOV2(V416L, C450A)(-6Jα)-PMI-M3 | S3 | This study |
| pCold-T7-lin-NLS-lin-LOV2(V416L, C450A)(-7Jα)-PMI-M3 | S3 | This study |
| pCold-T7-lin-NLS-lin-LOV2(V416L, C450A)(-8Jα)-PMI-M3 | S3 | This study |
| pCold-T7-lin-NLS-lin-LOV2(V416L, I539E)(-6Jα)-PMI-M3 | S3 | This study |
| pCold-T7-lin-NLS-lin-LOV2(V416L, I539E)(-7Jα)-PMI-M3 | S3 | This study |
| pCold-T7-lin-NLS-lin-LOV2(V416L, I539E)(-8Jα)-PMI-M3 | S3 | This study |

**Table S4. List of plasmids for the PMI-truncated LOV2(V416L)-PMI expression library used in the *in vitro* binding assay.**

### Table S5

| Plasmid name | Figures | Source or reference |
|---|---|---|
| pCold-T7-lin-NLS-lin-LOV2(C450A)(-6Jα)-PMI(S2A) | S5 | This study |
| pCold-T7-lin-NLS-lin-LOV2(C450A)(-6Jα)-PMI(E5A) | 5, S5 | This study |
| pCold-T7-lin-NLS-lin-LOV2(C450A)(-6Jα)-PMI(W7A) | S5 | This study |
| pCold-T7-lin-NLS-lin-LOV2(C450A)(-6Jα)-PMI(L10A) | S5 | This study |
| pCold-T7-lin-NLS-lin-LOV2(C450A)(-6Jα)-PMI(T1L, L10A) | 5, S5 | This study |
| pCold-T7-lin-NLS-lin-LOV2(C450A)(-6Jα)-PMI(A4L, L10A) | S5 | This study |
| pCold-T7-lin-NLS-lin-LOV2(C450A)(-6Jα)-PMI(N8L, L10A) | S5 | This study |
| pCold-T7-lin-NLS-lin-LOV2(C450A)(-6Jα)-PMI(L10A, P12A) | S5 | This study |
| pCold-T7-lin-NLS-lin-LOV2(C450A)(-6Jα)-PMI(L10A, P12R) | S5 | This study |
| pCold-T7-lin-NLS-lin-LOV2(C450A)(-6Jα)-PMI(L10A, P12Q) | S5 | This study |
| pCold-T7-lin-NLS-lin-LOV2(C450A)(-6Jα)-PMI-M4 | 5, S5 | This study |
| pCold-T7-lin-NLS-lin-LOV2(C450A)(-6Jα)-PMI-M5 | 5, S5 | This study |
| pCold-T7-lin-NLS-lin-LOV2(C450A)(-6Jα)-p53(17-26) | S5 | This study |
| pCold-T7-lin-NLS-lin-LOV2(C450A)(-6Jα)-p53(18-26) | S5 | This study |
| pCold-T7-lin-NLS-lin-LOV2(C450A)(-6Jα)-p53(17-27) | 5, S5 | This study |
| pCold-T7-lin-NLS-lin-LOV2(C450A)(-6Jα)-p53(18-27) | S5 | This study |
| pCold-T7-lin-NLS-lin-LOV2(I539E)(-6Jα)-PMI(S2A) | S5 | This study |
| pCold-T7-lin-NLS-lin-LOV2(I539E)(-6Jα)-PMI(E5A) | 5, S5 | This study |
| pCold-T7-lin-NLS-lin-LOV2(I539E)(-6Jα)-PMI(W7A) | S5 | This study |
| pCold-T7-lin-NLS-lin-LOV2(I539E)(-6Jα)-PMI(L10A) | S5 | This study |
| pCold-T7-lin-NLS-lin-LOV2(I539E)(-6Jα)-PMI(T1L, L10A) | 5, S5 | This study |
| pCold-T7-lin-NLS-lin-LOV2(I539E)(-6Jα)-PMI(A4L, L10A) | S5 | This study |
| pCold-T7-lin-NLS-lin-LOV2(I539E)(-6Jα)-PMI(N8L, L10A) | S5 | This study |
| pCold-T7-lin-NLS-lin-LOV2(I539E)(-6Jα)-PMI(L10A, P12A) | S5 | This study |
| pCold-T7-lin-NLS-lin-LOV2(I539E)(-6Jα)-PMI(L10A, P12R) | S5 | This study |
| pCold-T7-lin-NLS-lin-LOV2(I539E)(-6Jα)-PMI(L10A, P12Q) | S5 | This study |
| pCold-T7-lin-NLS-lin-LOV2(I539E)(-6Jα)-PMI-M4 | 5, S5 | This study |
| pCold-T7-lin-NLS-lin-LOV2(I539E)(-6Jα)-PMI-M5 | 5, S5 | This study |
| pCold-T7-lin-NLS-lin-LOV2(I539E)(-6Jα)-p53(17-26) | S5 | This study |
| pCold-T7-lin-NLS-lin-LOV2(I539E)(-6Jα)-p53(18-26) | S5 | This study |
| pCold-T7-lin-NLS-lin-LOV2(I539E)(-6Jα)-p53(17-27) | 5, S5 | This study |

| | | |
|---|---|---|
| pCold-T7-lin-NLS-lin-LOV2(I539E)(-6Jα)-p53(18-27) | S5 | This study |
| pCold-T7-lin-NLS-lin-LOV2(C450A)(-7Jα)-PMI(S2A) | S5 | This study |
| pCold-T7-lin-NLS-lin-LOV2(C450A)(-7Jα)-PMI(E5A) | 5, S5 | This study |
| pCold-T7-lin-NLS-lin-LOV2(C450A)(-7Jα)-PMI(W7A) | S5 | This study |
| pCold-T7-lin-NLS-lin-LOV2(C450A)(-7Jα)-PMI(L10A) | S5 | This study |
| pCold-T7-lin-NLS-lin-LOV2(C450A)(-7Jα)-PMI(T1L, L10A) | S5 | This study |
| pCold-T7-lin-NLS-lin-LOV2(C450A)(-7Jα)-PMI(A4L, L10A) | S5 | This study |
| pCold-T7-lin-NLS-lin-LOV2(C450A)(-7Jα)-PMI(N8L, L10A) | S5 | This study |
| pCold-T7-lin-NLS-lin-LOV2(C450A)(-7Jα)-PMI(L10A, P12A) | S5 | This study |
| pCold-T7-lin-NLS-lin-LOV2(C450A)(-7Jα)-PMI(L10A, P12R) | S5 | This study |
| pCold-T7-lin-NLS-lin-LOV2(C450A)(-7Jα)-PMI(L10A, P12Q) | S5 | This study |
| pCold-T7-lin-NLS-lin-LOV2(C450A)(-7Jα)-PMI-M4 | S5 | This study |
| pCold-T7-lin-NLS-lin-LOV2(C450A)(-7Jα)-PMI-M5 | S5 | This study |
| pCold-T7-lin-NLS-lin-LOV2(C450A)(-7Jα)-p53(17-26) | S5 | This study |
| pCold-T7-lin-NLS-lin-LOV2(C450A)(-7Jα)-p53(18-26) | S5 | This study |
| pCold-T7-lin-NLS-lin-LOV2(C450A)(-7Jα)-p53(17-27) | S5 | This study |
| pCold-T7-lin-NLS-lin-LOV2(C450A)(-7Jα)-p53(18-27) | S5 | This study |
| pCold-T7-lin-NLS-lin-LOV2(I539E)(-7Jα)-PMI(S2A) | S5 | This study |
| pCold-T7-lin-NLS-lin-LOV2(I539E)(-7Jα)-PMI(E5A) | 5, S5 | This study |
| pCold-T7-lin-NLS-lin-LOV2(I539E)(-7Jα)-PMI(W7A) | S5 | This study |
| pCold-T7-lin-NLS-lin-LOV2(I539E)(-7Jα)-PMI(L10A) | S5 | This study |
| pCold-T7-lin-NLS-lin-LOV2(I539E)(-7Jα)-PMI(T1L, L10A) | S5 | This study |
| pCold-T7-lin-NLS-lin-LOV2(I539E)(-7Jα)-PMI(A4L, L10A) | S5 | This study |
| pCold-T7-lin-NLS-lin-LOV2(I539E)(-7Jα)-PMI(N8L, L10A) | S5 | This study |
| pCold-T7-lin-NLS-lin-LOV2(I539E)(-7Jα)-PMI(L10A, P12A) | S5 | This study |
| pCold-T7-lin-NLS-lin-LOV2(I539E)(-7Jα)-PMI(L10A, P12R) | S5 | This study |
| pCold-T7-lin-NLS-lin-LOV2(I539E)(-7Jα)-PMI(L10A, P12Q) | S5 | This study |
| pCold-T7-lin-NLS-lin-LOV2(I539E)(-7Jα)-PMI-M4 | S5 | This study |
| pCold-T7-lin-NLS-lin-LOV2(I539E)(-7Jα)-PMI-M5 | S5 | This study |
| pCold-T7-lin-NLS-lin-LOV2(I539E)(-7Jα)-p53(17-26) | S5 | This study |
| pCold-T7-lin-NLS-lin-LOV2(I539E)(-7Jα)-p53(18-26) | S5 | This study |
| pCold-T7-lin-NLS-lin-LOV2(I539E)(-7Jα)-p53(17-27) | S5 | This study |
| pCold-T7-lin-NLS-lin-LOV2(I539E)(-7Jα)-p53(18-27) | S5 | This study |

**Table S5. List of plasmids for the LOV2-PMI mutant expression library used in the *in vitro* binding assay.**

### Table S6

| | LOV2-PMI | | LOV2-PMI-M3 | |
|---|---|---|---|---|
| | WT | −6Jα | WT | −7Jα |
| LOV2 | 1 | 1 | 1 | 1 |
| $K^+$ | 37 | 34 | 37 | 37 |
| $Cl^-$ | 32 | 30 | 32 | 34 |
| Water molecule | 13520 | 12447 | 13378 | 13477 |
| Total number of atoms | 56664 | 52280 | 56116 | 56441 |
| Box size in $Å^3$ | 69.171× 89.981× 68.974 | 71.550× 76.658× 72.204 | 79.027× 69.660× 78.576 | 73.462× 72.019× 81.095 |

**Table S6. Detailed information of the MD simulation systems. The box size indicated is the value after pressure equilibration.**

**Movie 1**

Translocation of the Opto-MDMi (LOVTRAP) actuator to the nucleus. HeLa cells transiently expressing H1-mCherry, NTOM20-mNeonGreen-LOV2 localizer, and Zdk1-miRFP703-NLS actuator were cultured in a 4-well glass-bottom dish. Time-lapse imaging was performed using a spinning disk confocal microscope. The cells were repeatedly illuminated with blue light at 2-min intervals. Images were acquired every 5 seconds for a total imaging time of 10 min.

**Movie 2**

Transcriptional activation of p53 by Opto-MDMi (LOVTRAP). HCT116 cells harboring a p53 transcriptional reporter and stably expressing Opto-MDMi (LOVTRAP) fragments were cultured on a 4-well glass-bottom dish. Time-lapse imaging was performed using a wide-field microscope. The cells were continuously illuminated with blue light at 24 hours. Images were acquired every 15 min. Stacked images were aligned using the StackReg ImageJ plugin. Total imaging time = 30 hours.

**Movie 3**

Translocation of the screened LOV2-PMI modules to the plasma membrane. HeLa cells transiently expressing the MDM2(1-125)-mCherry-HRasCT and the LOV2-PMI modules fused with miRFP703 were cultured in a 4-well glass-bottom dish. Time-lapse imaging was performed using a spinning disk confocal microscope. The cells were repeatedly illuminated with blue light at 1 min intervals. Images were acquired every 5 seconds for a total imaging time of 5 min.

**Movie 4**

Predicted protein structures of the LOV2-PMI fragments. Light green color and magenta color indicate the LOV2 core and the PMI peptide, respectively. Important residues (Phe3, Trp7, and Leu10) in PMI/PMI-M3 for the interaction with MDM2/MDMX are highlighted using ball-and-stick models.

**Movie 5**

Simulated movement of the LOV2-PMI fragments by MD simulation. The color assignments are the same as in Movie4. The movie was created by visualizing the protein structure every 10 nanoseconds based on the MD simulation results covering the entire 1 microsecond.

**Movie 6**

Translocation of the screened LOV2-PMI mutant module to the plasma membrane. HeLa cells transiently expressing MDM2(1-125)-mCherry-HRasCT and LOV2-PMI mutant modules fused with miRFP703 were cultured in a 4-well glass-bottom dish. Time-lapse imaging was performed using a spinning disk confocal microscope. The cells were repeatedly illuminated with blue light at 1-min intervals. Images were acquired every 5 seconds for a total imaging time of 5 min.

**Movie 7**

Transcriptional activation of p53 by Opto-MDMi (LOV2-PMI). HCT116 cells harboring a p53 transcriptional reporter and stably expressing Opto-MDMi (LOV2-PMI) fragments were cultured on a 4-well glass-bottom dish. Time-lapse imaging was performed using a wide-field microscope. The cells were continuously illuminated with blue light at 24 hours. Images were acquired every 15 min. Stacked images were aligned using the StackReg ImageJ plugin. Total imaging time = 30 hours.

**Movie 8**

Transcriptional activation of p53 by Opto-MDMi. HCT116 cells harboring a p53 transcriptional reporter and stably expressing Opto-MDMi fragments were cultured in a 4-well glass-bottom dish. Time-lapse imaging was performed using a wide-field microscope. The cells were continuously illuminated with blue light at 24 hours. Images were acquired every 15 min. Stacked images were aligned using the StackReg ImageJ plugin. Total imaging time = 30 hours.
